# Odorant receptors mediating avoidance of toxic mustard oils in *Drosophila melanogaster* are expanded in herbivorous relatives

**DOI:** 10.1101/2024.10.08.617316

**Authors:** Teruyuki Matsunaga, Carolina E. Reisenman, Benjamin Goldman-Huertas, Srivarsha Rajshekar, Hiromu C. Suzuki, David Tadres, Joshua Wong, Matthieu Louis, Santiago R. Ramírez, Noah K. Whiteman

## Abstract

Plants release defense volatile compounds that can deter herbivores. Among them are electrophilic toxins, such as isothiocyanates from mustard plants, that activate pain receptors by contact (i.e. taste) in many animals, including *Drosophila melanogaster*. While specialist insects have evolved strategies to tolerate toxicity and use mustard plants as hosts, it is unclear whether non-specialist insects detect and avoid electrophilic toxins via olfaction. To address this, and to understand if specialized insects co-opted these toxic compounds as hostplant olfactory cues, we leveraged closely related drosophilid species, including the microbe-feeding *D. melanogaster* and *Scaptomyza pallida*, and the mustard-feeding specialist *S. flava*. In olfactory assays, *D. melanogaster* exposed to allyl isothiocyanate volatiles were rapidly immobilized, demonstrating the high toxicity of this wasabi-derived compound to non-specialists. Through single sensillum electrophysiological recordings from olfactory organs and behavioral assays, we identified an Olfactory receptor (Or) necessary for volatile detection and behavioral aversion to allyl isothiocyanate in *D. melanogaster*. RNA sequencing and heterologous expression revealed that *S. flava* possess lineage-specific, triplicated homologs of this *Or,* and that each paralog exhibited broadened and distinct sensitivity to isothiocyanate compounds. Using AlphaFold2 modeling, site-directed mutagenesis and electrophysiological recordings, we identified two critical amino acid substitutions that changed the sensitivity of these paralogs from fruit-derived odors to isothiocyanates in the mustard specialist *S. flava*. Our findings show that non-specialists can detect electrophiles via olfaction, and that their olfactory systems can rapidly adapt to toxic hostplant niches through co-option and duplication of ancestral chemosensory genes with few amino acid changes.

## 1. INTRODUCTION

Plants have evolved the ability to synthesize a diverse array of toxic specialized metabolites that can provide resistance against insect herbivory. In turn, herbivorous insects have evolved diverse morphological, physiological, and behavioral counter-strategies to avoid these chemicals if encountered, or to mitigate their effects if ingested (Mithöfer and Boland 2012). Some herbivores even co-opt these plant toxins as oviposition or feeding stimulants (and even as chemical defenses of their own). For instance, monarch butterfly larvae evolved insensitivity against cardenolides released from their milkweed host plants (Reichstein et al. 1968, Dobler et al. 2012). Many plant toxins are, however, far more promiscuous in their modes of action, which presents a different “evolutionary hurdle” (Southwood 1972) to herbivores. Among them are various alkaloids, terpenoids, and electrophilic green leaf volatiles (Noge and Becerra 2015; Yaffe et al. 2015; Iorio et al. 2022) that intoxicate and deter herbivores by forming covalent bonds with biological molecules (War et al. 2012).

Mustard plants (Brassicales: Brassicaceae) such as thale cress (*Arabidopsis thaliana*), arugula (*Eruca sativa*), and wasabi (*Eutrema japonicum*) have evolved a sophisticated chemical defense system that produces electrophilic toxins (Ahuja et al. 2010). These plants produce non-toxic glucosinolates, some of which are hydrolyzed *in planta* to form toxic electrophilic compounds such as isothiocyanates (ITCs) (Hopkins et al. 2009). ITCs are reactive compounds defined by a −N=C=S functional group attached to the rest of the molecule, wherein the electron-deficient carbon is attacked by nucleophiles. Examples of ITCs include allyl ITC (AITC) derived from wasabi and radish *Raphanus sativus* (Cuellar-Nuñez et al. 2022), and butyl ITC derived from the cabbage *Brassica oleracea* (MacLeod et al. 1989). The chemical diversity of glucosinolates allows Brassicales plants to effectively deter a wide array of herbivorous insects because different species have different mixtures of glucosinolates, making it more difficult for insects to adapt (Winde and Wittstock 2011).

One herbivorous insect lineage that has evolved to cope with these toxic Brassicales metabolites are obligated leaf-mining drosophilid flies in the genus *Scaptomyza* (e.g., *S. flava* and *S. montana*), which is phylogenetically nested within the paraphyletic *Drosophila* subgenus. These *Scaptomyza* mustard specialists, through rapid gene duplication and non-synonymous changes, have evolved some of the most efficient ITC-detoxifying enzymes known from animals (Gloss et al. 2014, 2019).

While detoxification mechanisms help animals cope with at least some noxious compounds, sensory detection and behavioral avoidance of these substances can act as a checkpoint to prevent intoxication. Indeed, insects can behaviorally avoid toxic chemicals through gustation and/or olfaction (e.g. Bernays and Chapman 1987; Stensmyr et al. 2012; Scott 2018; Chen and Dahanukar 2020; Dweck and Carlson 2020). This includes avoidance of ITCs: exposure to volatile mustard plant extracts kills *D. melanogaster* (Lichtenstein et al. 1964), and physical contact with AITC triggers repulsion via gustatory receptor cells that express the nociceptive “wasabi receptor” TrpA1 and Painless (Al-Anzi et al. 2006; Kang et al. 2010; Kim et al. 2010; Mandel et al. 2018). Additionally, volatile AITC causes behavioral aversion in fire ants (*Solenopsis invicta*) (Hashimoto et al. 2019). However, the functional and genetic basis underlying olfactory detection and avoidance of electrophilic toxins like ITC remain poorly understood.

*Scaptomyza* species include both non-herbivorous (e.g., microbe-feeding) and herbivorous species that use Brassicales plants (Aguilar et al. 2024). Herbivorous species have lost olfactory receptors that ancestral microbe-feeding species (Whiteman et al. 2011, 2012; Peláez et al. 2023) like *D. melanogaster* use to detect fermentation, microbial, and fruit odors (Goldman-Huertas et al. 2015). Likely to aid hostplant location, the herbivorous specialist *S. flava* has evolved paralogous copies of the Olfactory receptor 67b (which is expressed in antennal olfactory sensory neurons, OSNs) that respond to ITCs. In contrast, the single copy of microbe-feeding *D. melanogaster* and *S. pallida* respond to green leaf volatiles like *trans*-3-hexenol but not to ITCs (Matsunaga et al. 2021). Although these findings provide insight into how sensory receptors evolved in specialists, how evolutionary changes facilitate aversion and/or attraction to toxic compounds is still largely unknown, yet it is a central problem in understanding how organisms invade toxic niches and herbivorous insects specialize on toxic host plants.

In this study, we addressed the following three questions (Figure 1): (1) Do non-specialist insects detect toxic volatile ITCs via olfaction and behaviorally avoid them? (2) Have the homolog olfactory receptors of specialist insects evolved broader sensitivity to ITC hostplant compounds? (3) If so, what molecular changes in these homologous olfactory receptors underlie this broadened sensitivity? To address these questions, we first conducted behavioral toxicity assays and found that *D. melanogaster* is rapidly immobilized by volatile exposure to AITC. We demonstrated that olfactory detection and behavioral avoidance of this compound requires Olfactory receptor 42a (Or42a), which is expressed specifically in maxillary palp OSNs (Ray et al. 2007). Next, in the specialist species *S. flava*, we discovered that the *Or42a* paralog is triplicated and that the number of Or42a-positive OSNs is expanded. Concomitantly, we found that these paralogous Or42a proteins are collectively sensitive to a broad range of ITC compounds, whereas the single Or42a copies encoded in the genomes of the microbe-feeding *S. pallida* and *D. melanogaster* respond only to AITC. Finally, AlphaFold2 3D modeling, site-directed mutagenesis, and electrophysiological experiments identified two key amino acid replacements that shifted the sensitivity of *S. flava* Or42a paralogs from fruit odors to ITCs. Collectively, our findings demonstrate that plant-derived volatile toxins like ITCs negatively impact non-specialists and are detected via olfaction to mediate avoidance, and that gene duplication events and tuning shifts of olfactory receptors are coupled with specialization of herbivorous insects onto toxic hostplants.

**Figure 1:**
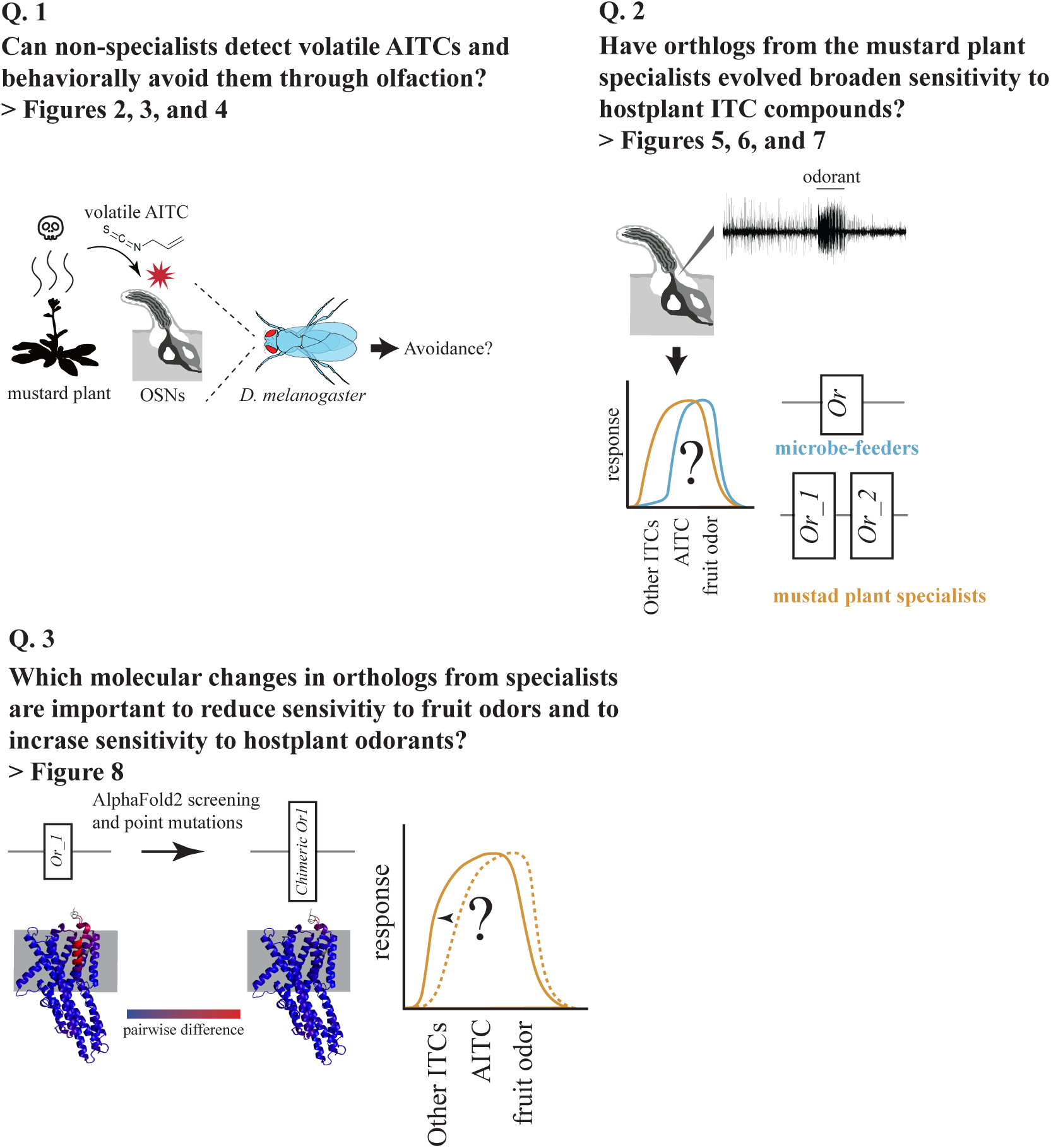
Questions addressed in this study.

## 2. RESULTS

### 2-1. Volatile allyl isothiocyanate (AITC) rapidly immobilizes *D. melanogaster*

To assess the toxic effects of volatile AITC in *D. melanogaster*, we conducted immobility assays with various concentrations of volatile AITC (from 1:500 to 1:2.5 vol/vol). In our experimental setup, a fabric mesh separated the chamber containing the flies from the chamber containing the AITC solution and therefore, flies were exposed to volatile AITC but could not contact (i.e. taste) the AITC solution directly (Figure Supplement 1A). While all insects in both the control treatment and those exposed to AITC 1:250 and 1:500 vol/vol remained active, most flies exposed to AITC concentrations ≥ 1:50 vol/vol became paralyzed (likely highly intoxicated or even dead) within 10 minutes (Figure 2). Thus, this rapid immobilization indicates that plant-derived electrophilic compounds like AITC can have a strong negative effect on flies, consistent with previous findings (Lichtenstein et al. 1964).

**Figure 2:**
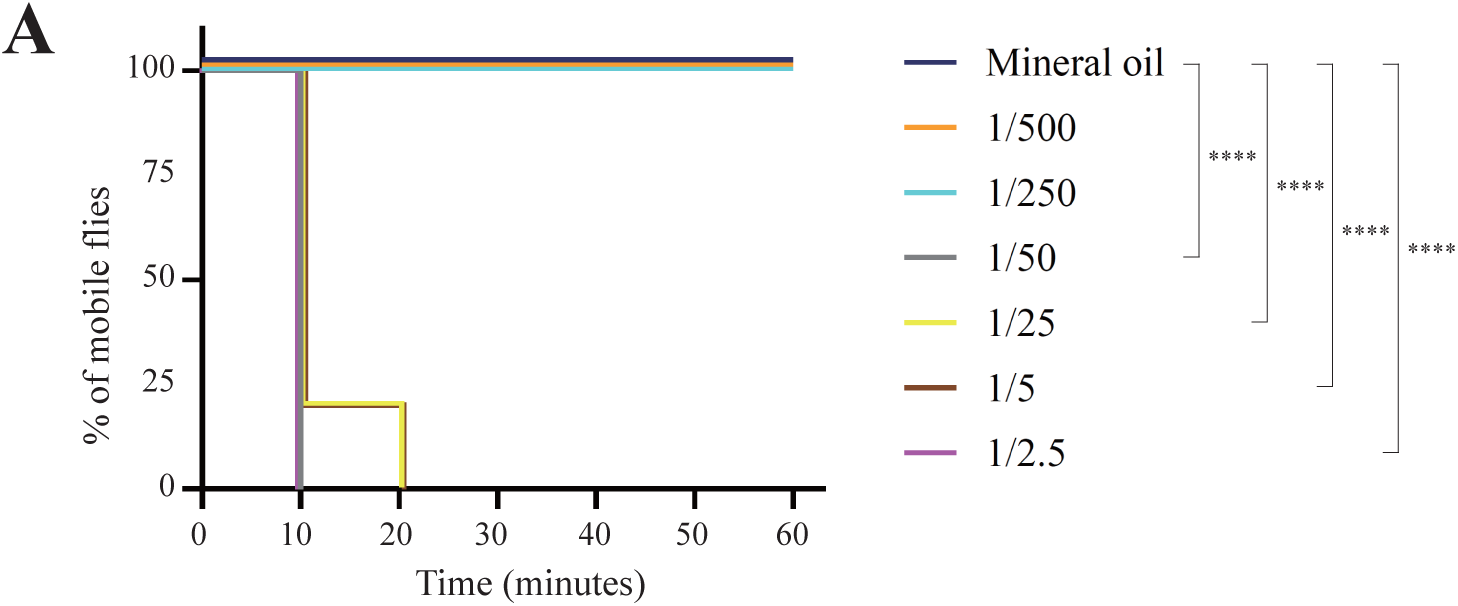
*Drosophila melanogaster* is immobilized rapidly upon exposure to volatile allyl isothiocyanate. The toxic effect of various concentrations (vol/vol) of volatile AITC to *D. melanogaster* was assessed by measuring the % of mobile flies; flies were not allowed to contact the AITC source (Figure Supplement 1A). Increasing the concentration of AITC decreases the % of mobile flies, likely due to intoxication. n=10 for each condition (solvent and concentration). **** p<0.0001, log-rank Mantel-Cox tests against the control.

### 2-2. Volatile AITC is detected by the maxillary palp olfactory receptor Or42a in *D. melanogaster*

The high toxicity caused by volatile AITC indicates that this volatile had the potential to be detected by the fly’s olfactory system, and that this could provide a fitness benefit if these volatile toxins were then behaviorally avoided. We investigated this in *D. melanogaster* by conducting exhaustive single sensillum recordings (SSR) from the fly’s olfactory organs, the antennae and the maxillary palps, upon stimulation with volatile AITC. Several olfactory sensory neurons (OSNs) showed excitatory responses to AITC, but OSNs in palp basiconic sensilla 1a (pb1a) were the most activated (Figure 3A-B, >100 spikes/sec). Because pb1a OSNs express Or42a (Couto et al. 2005), we investigated whether OSN responses in this sensilla type are indeed mediated by this olfactory receptor. OSNs in pb1 sensilla from genetic background control flies (*w1118*) showed strong responses to volatile AITC, while these OSNs showed no response in *Or42a* -/- flies (Figure 3C, <2 spikes/second), indicating that Or42a detects volatile AITC in these OSNs. Furthermore, OSNs in pb1a of *TrpA1^1^* null mutant flies responded strongly to AITC (>150 spikes/second, Figure 3C), showing that the contact chemoreceptor TrpA1 is not necessary for maxillary palp olfactory detection of this compound.

**Figure 3:**
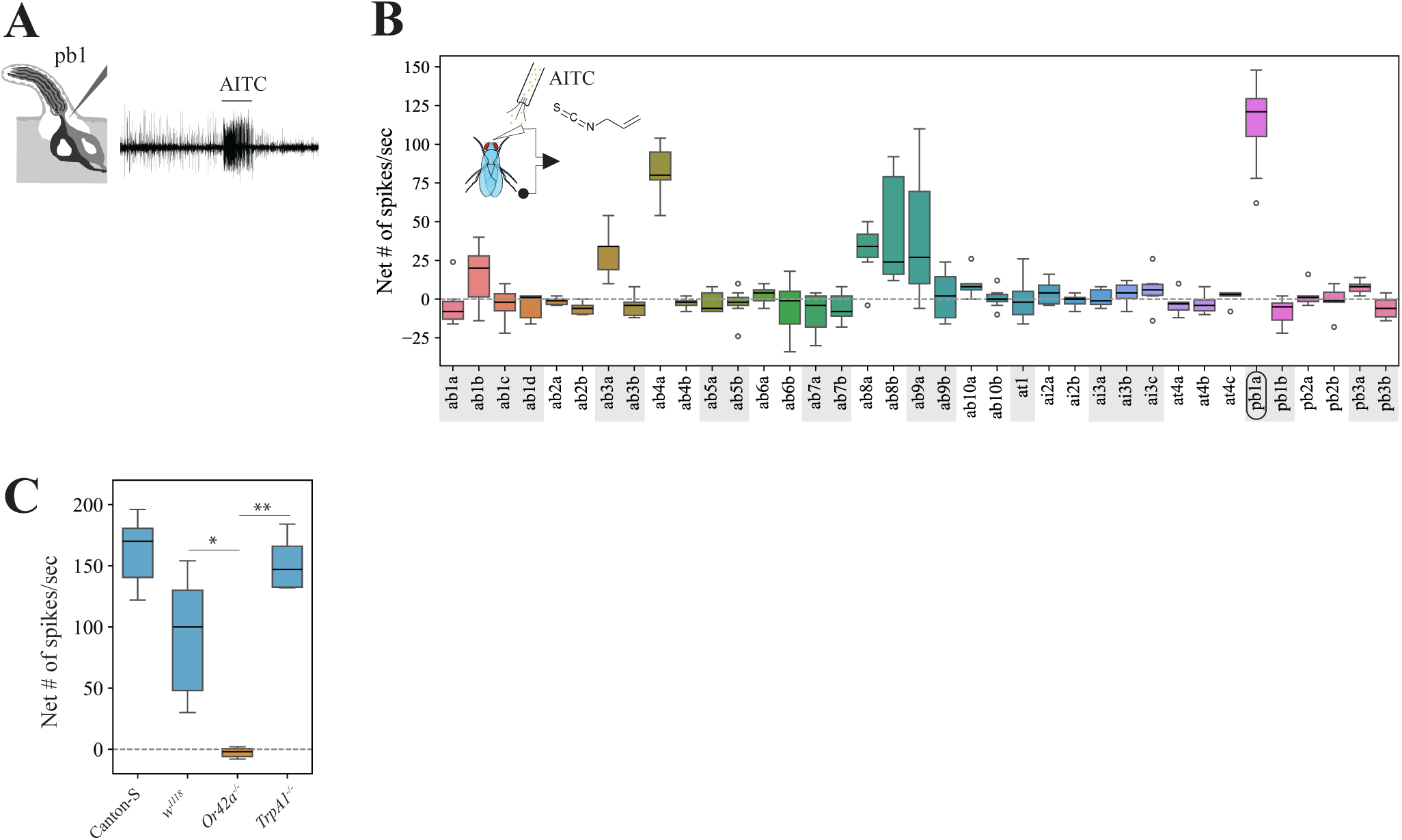
*Drosophila melanogaster* Or42a mediates detection of volatile allyl isothiocyanate. **(A)** Schematic and representative trace of a single sensillum recording (SSR) from pb1 OSNs upon stimulation with AITC 1:100 vol/vol. The horizontal bar indicates the onset of the stimulus and its duration (1 sec). **(B)** SSR from all *D. melanogaster* antennae and palp basiconic sensilla upon AITC stimulation (1:100 vol/vol; n= 6-10 recordings/sensilla type from 6 animals). Represented here and in all figures are the control-subtracted net number of spikes/sec, unless otherwise noted. The horizontal dotted line at zero indicates no response to odor stimulation. Here and thereafter, horizontal bars represent the median, the edges of the boxes correspond to 25^th^ and 75^th^ quartiles, the whiskers denote 10^th^ and 90^th^ quartiles, and symbols indicate outliers. Pb1a sensilla, which house Or42a (Ray et al. 2007), respond strongly to AITC. **(C)** Responses from pb1 sensilla of wildtype Canton-S, the genetic background control *w^1118^*, *Or42a-/-*, and *TrpA1^1^ D. melanogaster* flies to 1:100 vol/vol AITC stimulation (n=6-10 sensilla/genotype from 3-4 animals/genotype). The responses of both mutant flies were compared against each other and against those of *w1118*. Kruskal-Wallis ANOVA followed by Dunn’s multiple comparisons, *p<0.05, **p<0.01.

### 2-3. Volatile allyl isothiocyanate repels *D. melanogaster* via the olfactory receptor Or42a

Next, we examined if Or42a plays a role at the behavioral level. We conducted a positional olfactory assay based on Ohashi and Sakai (2015) with modifications to prevent flies from physically contacting the odor source. Female flies (n=10-12) were released in a dispositive consisting of two glass tubes, each of which was connected to a vial containing an odor solution or a vial loaded with a solvent (Figure Supplement 1B). The number of flies in each tube (hereafter “odorless/control tube” and “odorous/test tube”) as well as in the release section was counted every five minutes up to minute 35 (and again at 65 minutes). To validate this assay, we confirmed that wildtype *D. melanogaster* flies (strain Canton-S) showed normal olfactory-guided behavior in this behavioral set-up using apple cider vinegar (one-sample signed rank tests, p<0.05 in all cases, n=21, Supplementary File 4), a well-established *D. melanogaster* olfactory attractant (Semmelhack and Wang 2009; Becher et al. 2010). We then tested whether AITC causes olfactory repellence. For these and all forthcoming behavioral experiments, we used as a genetic background control an “empty Gal4 control” line (line # 68384) which carries *white* in the background of a *white* mutation, as in all the mutant fly lines. These control flies and wild-type Canton-S flies avoided the AITC tube at various time points (25, 30, 35 and 65 minutes, Figure 4A and Figure Supplement 2A). In contrast, *Or42a-/-* mutants avoided the AITC tube only at the 65 minutes time point (Figure 4A), possibly by taste detection of AITC via TrpA1 (volatile AITC molecules might have adhered to the glass tube walls at this point). Importantly, we confirmed that *Or42a-/-* mutants are capable of odor-mediated olfactory orientation in dual-choice trap assays offering apple cider vinegar vs. water (described in Matsunaga et al., 2021): all captured flies were recruited to the odor-baited trap (N=8 tests, n=20 females/test; Wilcoxon-matched pairs test, p<0.005; Supplementary File 4). These results suggest that *Or42a* plays a crucial role in mediating olfactory-driven behavioral aversion to AITC.

**Figure 4:**
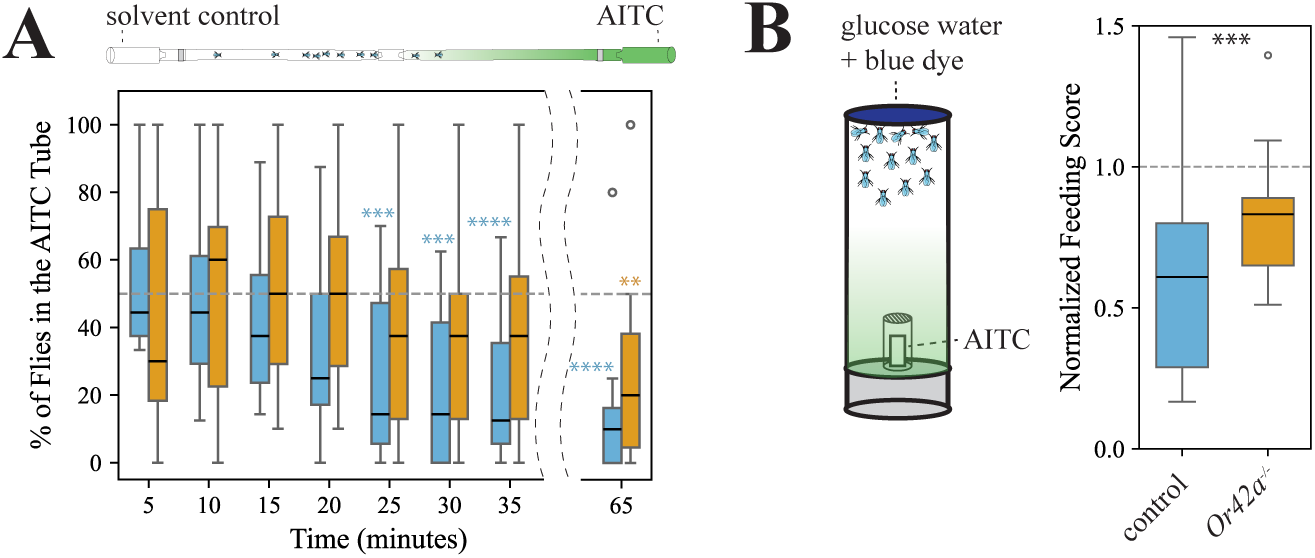
*Drosophila melanogaster* Or42a mediates behavioral aversion to allyl isothiocyanate. **(A)** Positional olfactory assay. Flies could smell, but not contact, the AITC solution (Figure Supplement 1B). The number of flies in the odorless and the odorous glass tubes was counted every five 5 minutes until minute 35, and then again at 65 minutes. The dotted line at 50% indicates random distribution between the two tubes. Genetic background control flies (line # 68384, blue boxes, n=15) avoided the tube closest to the odor source at various time points (***p<0.005, ****p<0.001; one-sample signed rank tests against median=50%). *Or42a-/-* mutants (orange boxes, n =15) distributed randomly between the two tubes at all timepoints (p>0.05) except at 65 minutes (*p<0.01). **(B)** Food consumption assay in presence or absence of AITC volatiles (1:500 vol/vol). Flies could smell but not contact the AITC solution (Figure Supplement 1C). Both genetic background control flies (n=16) and *Or42a -/-* mutants (n=14) fed less in the presence of AITC volatiles (medians<50%, one-sample signed rank tests on normalized data; respectively p<0.05 and p<0.001), but the feeding score of genetic control flies was lower than that of mutant flies (Mann-Whitney U test, ***p<0.001). The dotted line at 50% indicates no feeding aversion or enhancement.

Given that detection of food-related volatiles is known to increase sugar consumption (Reisenman and Scott, 2019), and that palp OSNs are located near the mouthparts, we reasoned that volatile detection of toxic/aversive odors such as AITC would instead suppress sugar consumption. In each test, we offered starved *D. melanogaster* (n=10-15 females/test) 50 mM glucose water solution dyed blue for 15 minutes in presence or absence of an odor: one group of flies was exposed to volatile AITC (1:500 vol/vol; flies could not contact the odor solution), and the control group was exposed to the mineral oil solvent (Figure Supplement 1C; flies could not contact the odor source). We calculated a feeding score/test (each vial constitutes a biological replicate) based on the amount of blue dye in the abdomen of flies. Control flies fed less when volatile AITC was present than in the presence of the solvent (one-sample signed rank tests on normalized data, p<0.001; Figure 4B). *Or42a-/-* mutant flies fed less in the presence of AITC volatiles as well (p<0.05), but their feeding scores were higher than those of the control group (p<0.001, Mann-Whitney U test; Figure 4B). Altogether, these results show that Or42a mediates aversion to AITC volatiles in two different behavioral contexts.

While we showed that Or42a-positive OSNs mediate behavioral aversion to AITC (Figure 4), Dweck et al. (2016) found that these OSNs mediate attraction to some fruit/fermentation volatiles. We thus tested whether Or42a also mediates behavioral attraction to such compounds in our experimental set-up/s using γ-hexalactone, which has been reported to activate Or42a-positive OSNs (Dweck et al., 2016). In the positional olfactory assay, genetic background control and wildtype Canton-S flies, but not *Or42a-/-* mutants, were attracted γ-hexalactone 1:10 vol/vol at various time points (one-sample signed rank tests, p<0.05 for both lines; Figure Supplement 2D and Supplementary File 4). Similarly, in consumption assays, Canton-S flies increased their feeding in the presence of volatile γ-hexalactone 1:50 vol/vol, but this effect was lost in *Or42a-/-* mutants (Figure Supplement 2J). Additionally, we found that volatile γ-hexalactone, contrary to what we observed in tests with AITC, does not immobilize flies (Figure Supplement 2K). Thus, our behavioral assays show that volatile γ-hexalactone attracts *D. melanogaster* via *Or42a* and is harmless, in line with previous results (Dweck et al., 2016).

Given that Or42a OSNs mediates both repellence to AITC and attraction to γ-hexalactone (Figure 4 and Figure Supplement 2A-E, J), we hypothesized that additional OSNs contribute to mediate these contrasting behavioral responses. We performed exhaustive SSR from antennae and maxillary palps and found that pb1a OSNs were the only ones activated by γ-hexalactone among basiconic, intermediate, and trichoid sensilla (Figure Supplement 2L). Due to technical limitations, we were unable to study antennal coeloconic sensilla, but Oh et al. (2021) reported that this odor activates Or35a OSNs in ac3b (Yao et al. 2005). In contrast, AITC activated several OSNs within the above mentioned sensilla types, including Or7a OSNs (>75 spikes/second, Figure 3B), which are housed in ab4a (Lin et al. 2015). Thus, we investigated whether *Or7a* and *Or35a* could respectively contribute to mediate the observed behavioral repellence to AITC and attraction to γ-hexalactone. Compared to genetic background controls, *Or7a -/-* flies showed less aversion to AITC in the positional olfactory assay (Figure Supplement 2F-G), while *Or35a -/-* flies lost the attraction to γ-hexalactone (Figure Supplement 2H-I). Thus, not only Or42a, but Or35a and Or7a are also necessary to mediate attraction/repellence behaviors to these odorants: activation of both Or42a and Or7a OSNs by AITC might mediate aversion, while activation of both Or42a and Or35a OSNs by γ-hexalactone might mediate attraction. Our findings thus support a combinatorial hypothesis of odor coding with respect to these ligands, and indicate that the valence of Or42a-mediated behavioral responses is context-dependent (Figure Supplement 2M).

### 2-4. Pb1a-like OSNs in Brassicales-specialists evolved broadened sensitivity to isothiocyanates

We next investigated how the evolutionary transitions from microbe-feeding to herbivory have affected the odor tuning of OSNs, using the reference species *D. melanogaster* as well as species within *Scaptomyza*: the microbe-feeding *S. hsui* and *S. pallida*, and the herbivorous Brassicales specialists *S. flava* and *S. montana* (Kim et al. 2021; Peláez et al. 2023). We investigated whether *Scaptomyza* species have pb1-like sensilla homologous to *D. melanogaster* pb1 and if so, the extent to which they respond to a broader range of ITC compounds, since Brassicales plants release many different ITCs upon wounding (MacLeod et al. 1989; Cuellar-Nuñez et al. 2022).

We first validated our methods for functional characterization of sensilla. Using compounds that serve as diagnostic for the three sensilla types found in the maxillary palps of *D. melanogaster* (see methods for details), various other plant-derived volatiles and several Brassicales-derived ITCs, we confirmed the presence of three different types of palp sensilla (pb1, pb2 and pb3) in this species (Figure 5 and Figure Supplements 3-4). As we observed before, *D. melanogaster* pb1a OSNs responded to AITC (>120 spikes/sec), but not to the other ITCs tested (<10 spikes/sec, Figure 5). We next characterized OSNs in pb sensilla in the four *Scaptomyza* species and used the spike rate to determine if data clustered by sensilla type (Figure 5, Figure Supplement 3-4). In all *Scaptomyza* species sensilla fell into three functional classes, two of which were functionally similar to *D. melanogaster* pb1 and pb2 (we termed them pb1-like and pb2-like). The third class was functionally different to any *D. melanogaster* palp sensilla type and clustered separately (termed pb3-like, Figure Supplement 3-4). As in *D. melanogaster*, AITC activated pb1a-like sensilla in all four *Scaptomyza* species (Figure 5) but several other ITCs, including isobutyl ITC, butyl ITC, and *sec*-butyl ITC, additionally activated *S. flava* and *S. montana* pb1a-like OSNs (Figure 5).

**Figure 5:**
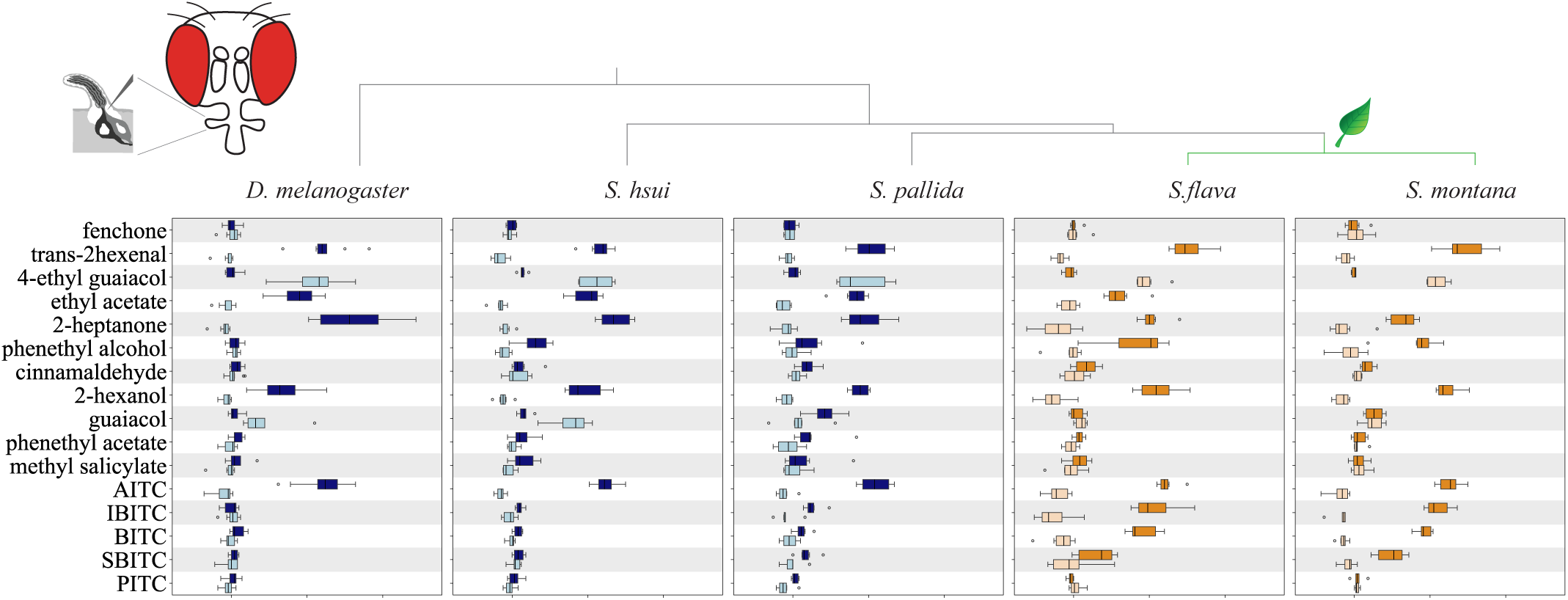
Maxillary palp pb1-like sensilla of the mustard plant specialist *Scaptomyza* species have an expanded ITC-sensitivity range. Single sensillum recordings from maxillary palp pb1 OSNs of *D. melanogaster, S. hsui, S. pallida, S. flava,* and *S. montana*. Stimuli (1:100 vol/vol) included diagnostic chemicals used to identify Ors in *D. melanogaster* (see Methods), fruit volatiles, green leaf volatiles, and Brassicales plant-derived isothiocyanates (ITCs) (n=6-9 from 3-4 animals/species). Pb1 and pb1-like sensilla housed two OSNs, labeled “a” (darker color) and “b” (lighter color). See methods for sensilla classification and the Figure supplement 3-4 for functional characterization of pb2 and pb3. While pb1a OSNs from all species responded to AITC, pb1a-like OSNs from the mustard plant specialists *S. flava* and *S. montana* additionally responded to other ITC compounds. Mustard specialization occurred at the clade leading to the common ancestor of these two species, denoted by the leaf cartoon. Because sensilla with extremely small spike amplitudes were excluded from analysis, additional unidentified palp basiconic sensilla may exist in *Scaptomyza*. AITC: allyl ITC, IBITC: isobutyl ITC, BITC: butyl ITC, SBITC: *sec*-butyl ITC, PITC: phenethyl ITC. The structure of ITCs is shown in Figure Supplement 5.

We also conducted dose-responses to various odorants from pb1a/pb1a-like sensilla of the microbe-feeding species (*D. melanogaster, S. hsui* and *S. pallida*) and the Brassicales specialists (*S. lava* and *S. montana*). For the most part, responses increased with increasing odorant concentration (Figure Supplement 5A). For comparing odor sensitivity across species, we calculated the odorant concentration required to elicit a biological response halfway between the baseline and the maximum (50% effective concentration, EC_50_; Figure Supplement 5D). In agreement with its herbivore habit, the EC_50_ for the fruit odor γ-hexalactone was higher in the herbivore *S. flava* than in the microbe-feeding *S. hsui* (*S. flava* had even lower sensitivity to this odorant than the herbivore *S. montana*; Figure Supplement 5B and D). The EC_50_ for *trans*-hexenal was lower in the two herbivores species than in *D. melanogaster* or *S. hsui* (Figure Supplement 5D), and *S. flava* was even more sensitive to this odorant than *S. montana* (Figure Supplement 5C). Similarly, the EC_50_ for AITC was lower in all the *Scaptomyza* species, but the two mustard specialists had similar EC_50_ for all the ITCs tested (Figure Supplement 5D, right). Overall, all these findings suggest that microbe-feeding species exhibit higher sensitivity to the fruit odor γ-hexalactone, whereas Brassicales specialists show heightened sensitivity to electrophilic *trans*-2-hexenal and ITCs.

### 2-5. *Or42a* is triplicated in the genome of *S. flava* and is highly expressed in the maxillary palps

Genomic analysis across the *Scaptomyza* genus revealed a duplication of *Or42a* in the lineage leading to all known *Scaptomyza*, while the *D. melanogaster* outgroup had a single *Or42a* homolog (Figure Supplement 6A). Notably, syntenic analysis indicated that *S. flava* has a species-specific tandem triplication in one of the *Or42a* duplicates, resulting in three tandem paralogs, which we named *Or42a2*, *Or42a3*, and *Or42a4* (Figure 6A and Figure Supplement 6A-B). In contrast, *S. montana*, *S. graminum, S. pallida* and *S. hsui* retained only two *Or42a* paralogs.

**Figure 6:**
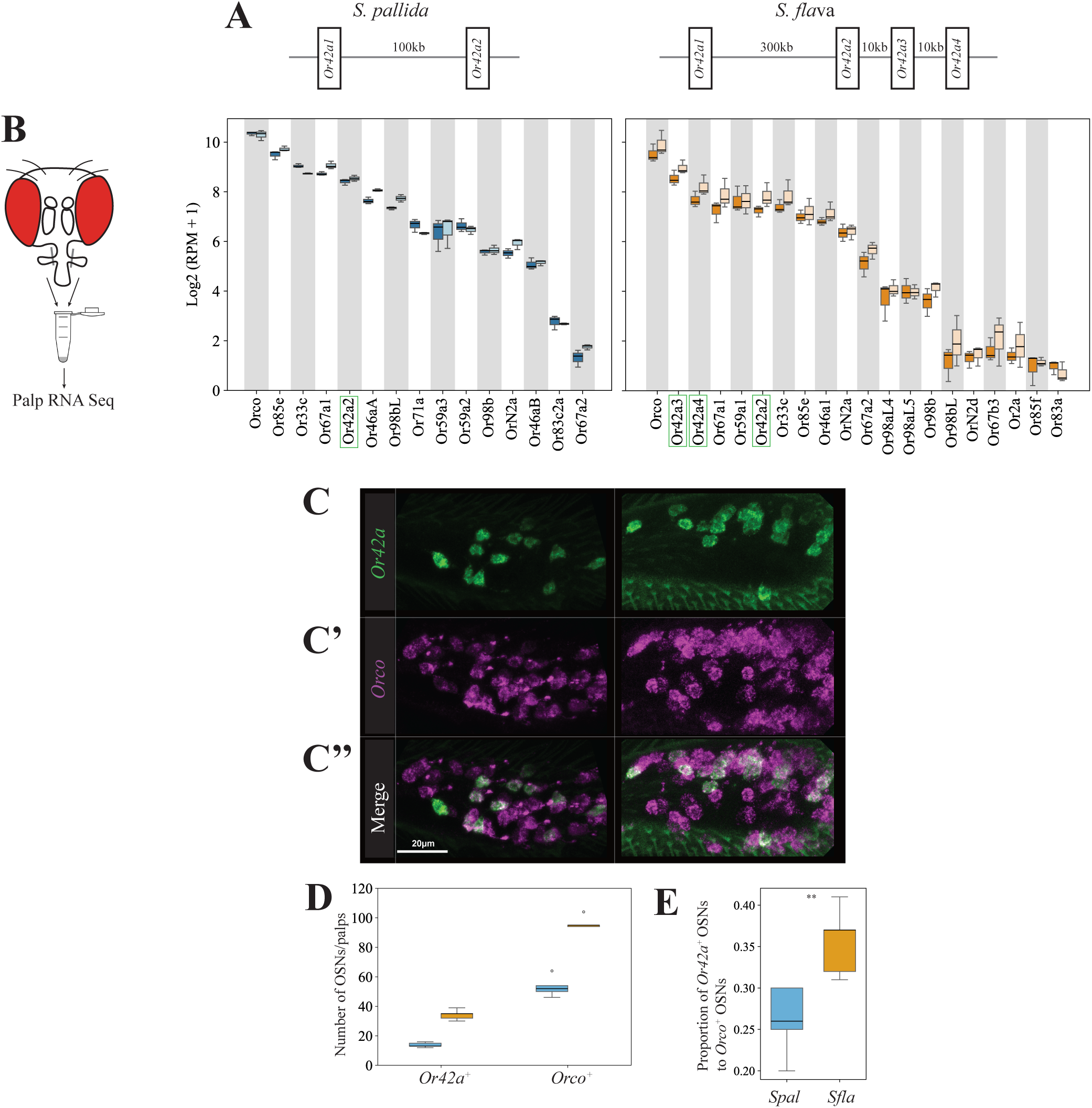
High expression of *Or42a* paralogs and over-representation of *Or42a*-positive OSNs in the maxillary palp of *S. flava*. **(A)** Schematic of *Or42a* syntenic regions in the genomes of the microbe-feeding *S. pallida* and the mustard plant specialist *S. flava*, with a gene triplication in the *S. flava* genome at the syntenic region of *S. pallida Or42a2* (*S. flava Or42a2*, *S. flava Or42a3*, and *S. flava Or42a4*). **(B)** Maxillary palp RNA seq of *S. pallida* and *S. flava Or*s (n=3 replicates/sex and species). *Or*s with median values of the log2 (RPM +1) < 1 (n=3) were excluded. *S. pallida* expresses only one copy of Or42a, while the specialist *S. flava* expresses three copies. **(C-C”)** Representative images of hybridization chain reaction RNA fluorescent in situ hybridization from the maxillary palps of *S. pallida* and *S. flava* showing *Or42a*-positive OSNs (**C**, green), *Orco*-positive OSNs (**C’**, magenta), and the merged signals (**C’’,** white indicates co-localization of *Or42a*-positive OSNs and *Orco*-positive OSNs). Or42a is expressed in OSNs. Scale bar: 20µm. **(D-E)** Number of *Or42a*-positive OSNs and *Orco*-positive OSNs in the maxillary palps of *S. pallida* (blue boxes) and *S. flava* (orange boxes) **(D),** and ratio between them **(E)**. Mann-Whitney U tests, ** p<0.01; n=5 animals/species. *S. flava* has more Or42a-positive OSNs and more Orco-positive OSNs than *S. pallida*.

Given these differences in the Or42a gene copy number across species, we conducted species- and sex-specific RNA transcriptome analyses of OSNs in the maxillary palps of the microbe-feeding *S. pallida* and the mustard plant specialist *S. flava*. We confirmed the expression of *S. pallida Or42a* and *S. flava Or42a2-4* in these organs (Figure 6B; Figure Supplement 7). Interestingly, the *S. flava Or42a* paralogs were each expressed at levels comparable to those of other *Or* genes, such as *Or33c*, the homolog of which is expressed in pb2a OSNs in *D. melanogaster* (Figure Supplement 7). We found that *S. pallida Or42a2* and *S. flava Or42a2-4* were expressed in the respective maxillary palp of each species, whereas *S. pallida Or42a1* and *S. flava Or42a1* were not expressed (Figure 6B).

### 2-6. The Brassicales specialist *S. flava* has more *Or42a*-positive olfactory sensory neurons

Given that pb1a-like OSNs likely express *Or42a* (Figure 6B), we hypothesized that the number of *Or42a*-positive OSNs is higher in the specialist species. To test this, we quantified the number of *Or42a*-positive OSNs in *S. flava* and *S. pallida* using hybridization chain reaction RNA fluorescent in situ hybridization (RNA FISH). We designed a *S. flava Or42a* RNA probe based on the conserved sequence region of *S. flava Or42a2*, *3*, and *4*, as the high sequence similarity among these paralogs prevented the design of paralog-specific probes. We found that the maxillary palps of *S. flava* contained more *Or42a*-positive OSNs and more *Orco*-positive OSNs (Orco is a highly conserved co-receptor necessary for odorant receptor olfactory function, Larsson et al., 2004) than those of *S. pallida* (Figure 6C-D). Because all maxillary palp OSNs express Orco (Larsson et al. 2004), the number of Orco-positive OSNs represents the total number of palp OSNs. Importantly, the ratio of the total number of *Or42a*-positive OSNs to the total number of *Orco*-positive OSNs was higher in *S. flava* (Figure 6E). These results indicate that *S. flava* has not only more *Or42a*-positive OSNs in the maxillary palps, but also a greater proportion of *Or42a*-positive OSNs relative to the overall number of OSNs. Accordingly, we predicted that the number of pb1-like sensilla would be also overrepresented in mustard specialists. To test this, we generated functional anatomical maps of sensilla on the anterior part of the maxillary palps of the *Scaptomyza* species (and of *D. melanogaster* for comparison) using diagnostic chemicals (Figure Supplement 8-9, Supplementary file 10). While *D. melanogaster* had a relatively randomized distribution of sensilla on the palps, consistent with previous reports (de Bruyne et al. 1999), all four *Scaptomyza* species exhibited a more organized sensilla pattern, with pb1-like, pb2-like, and pb3-like respectively located medially, distally, and proximally, as reported in *D. mojavensis*, which is more closely related to all *Scaptomyza* spp. than *D. melanogaster* (Crowley-Gall et al. 2016). We then quantified the number of each sensilla type across species and found that both *S. flava* and *S. montana* have a larger number of pb1-like than of pb2-like or pb3-like sensilla, while *D. melanogaster* and the other two microbe-feeding *Scaptomyza* species had similar proportions of each sensilla type (Figure Supplement 8-9). These findings, showing a triplication of Or42a, along with an expansion in the number of Or42a-positive OSNs and pb1a-like sensilla, are in line with the enhanced capacity of Brassicales-specialist *Scaptomyza* species to detect volatile ITCs.

### 2-7. Paralog-specific functional evolution of the olfactory receptor Or42a

We next investigated whether the increased sensitivity of *S. flava* pb1a-like OSNs to ITCs (Figure 5) also resulted from changes in the odor tuning of the Or42a triplicates. To test this, we expressed *D. melanogaster* Or42a*, S. pallida* Or42a2, and *S. flava* Or42a2-4 in *D. melanogaster* antennal trichoid 1 (at1) OSNs (in the background of a null mutation for *Or67d*, the at1 cognate receptor Kurtovic et al. 2007) and conducted functional analysis. We found that γ-hexalactone, *trans*-2-hexenal, and AITC strongly activated OSNs expressing *D. melanogaster* Or42a (>79 spikes/second) or *S. pallida* Or42a2 (>96 spikes/sec), while the other ITC compounds evoked much weaker responses from these two orthologs (respectively <11 and 18 spikes/second; Figure 7A-B). OSNs expressing *S. flava* Or42a3 or *S. flava* Or42a4 were also very sensitive to AITC and *trans*-2-hexenal but additionally responded to isobutyl ITC and butyl ITC (> 87 and 46 spikes/second, respectively; Figure 7B), consistently with the odor response profiles of *S. flava* pb1a OSNs (Figure 5). Notably, OSNs expressing *S. flava* Or42a4 showed only small responses to γ-hexalactone (<10 spikes/second), and *S. flava* Or42a2 was only activated by AITC (Figure 7B). These results are in line with the broader ITC sensitivity of *S. flava* pb1a OSNs compared to that of *D. melanogaster* pb1a and *S. pallida* pb1a-like OSNs. Altogether, our findings reveal paralog-specific functional evolution of Or42a triplicates in *S. flava*, wherein different paralogs evolved distinct sensitivities to different ITCs and fruit odors.

**Figure 7:**
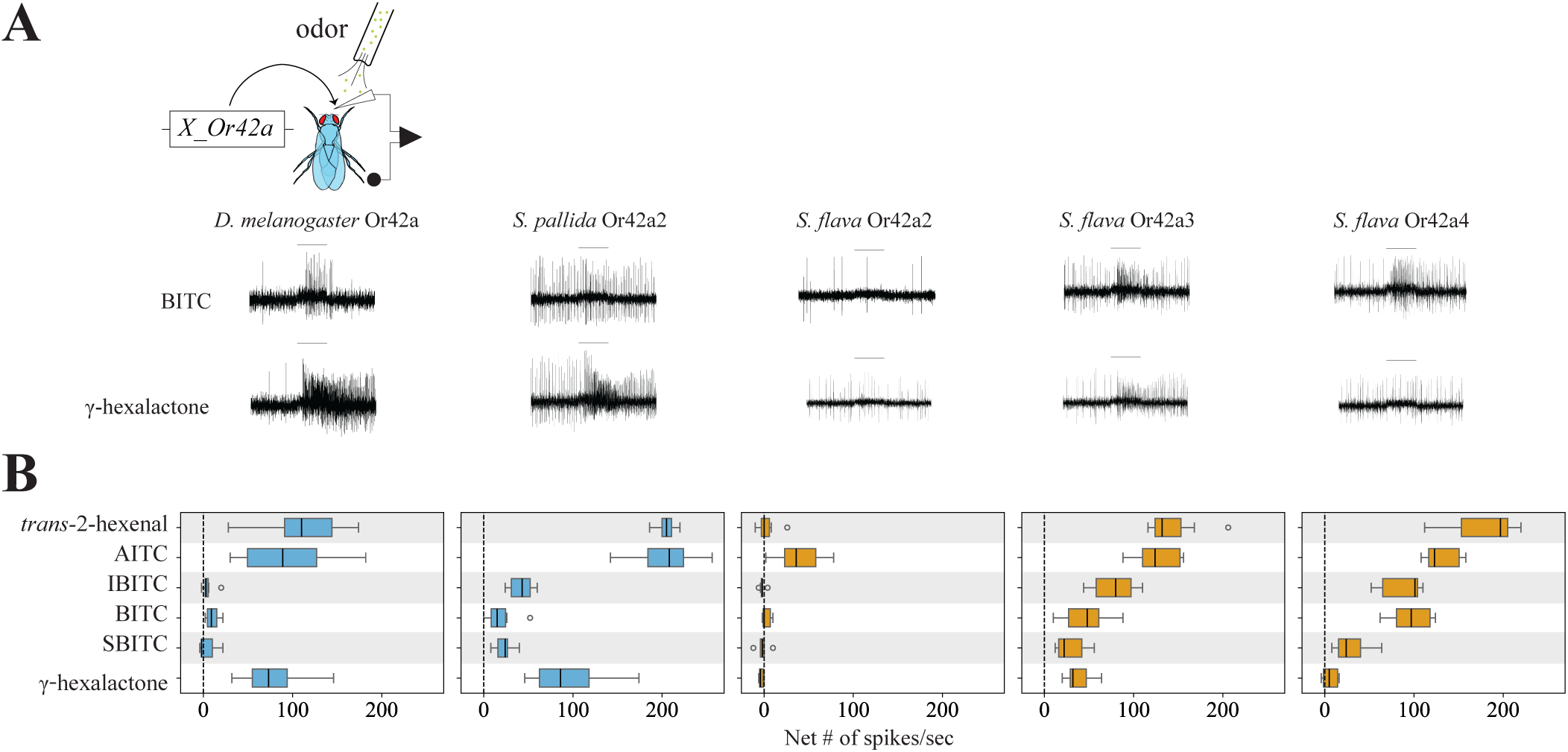
Functional characterization of the olfactory receptor Or42a from *D. melanogaster*, *S. pallida*, and *S. flava.* **(A)** Representative SSR traces from *D. melanogaster* at1 sensilla OSNs expressing species-specific *Or42a* under the control of *Or67d ^Gal4^* (fly genotype: *UAS-Or42a; Or67d ^Gal4^*) in response to stimulation with butyl isothiocyanate (BITC) and the fruit odor γ-hexalactone. The horizontal bars above records indicate the onset and duration (1 sec) of the stimulation. **(B)** Responses of at1 OSNs (n=6-8 sensilla from 3-4 animals/genotype) expressing species-specific *Or42a* upon stimulation with *trans*-2-hexenal (a general leaf odor released upon leaf mechanical damage such as crushing), various ITCs produced by mustard plants (AITC, IBITC, BITC, SBITC, abbreviations as in Figure 5), and γ-hexalactone. Orthologs from all species respond to AITC, while *S. flava* Or42a3 and *S. flava* Or42a4 additionally respond to various ITCs. In contrast, only paralogs from microbe feeding species show strong responses to γ-hexalactone.

### 2-8. AlphaFold2-led screening with ectopic expression of *S. flava* Or42a reveals the molecular changes underlying changes in odor sensitivity

We next investigated which amino acid substitutions in *S. flava* O42a4 may have led to the gain of sensitivity to butyl ITC and the decreased sensitivity to γ-hexalactone (there are 32 amino acid differences between *S. flava* Or42a3 and *S. flava* Or42a4, Figure Supplement 10A). To explore the structural basis of these functional differences, we predicted the 3D structures of *S. flava* Or42a3 and *S. flava* Or42a4 and aligned the resulting models in 3D space using PyMol (Jumper et al. 2021; Mirdita et al. 2022; Benton and Himmel 2023; Himmel et al. 2023).

We first confirmed that the predicted local distance difference test scores for the *S. flava* Or42a3 and *S. flava* Or42a4 structures were sufficiently high to ensure confidence in the 3D predictions, except for the N-terminal and C-terminal regions (Figure Supplement 10B). The most striking 3D structure difference between *S. flava* Or42a3 and *S. flava* Or42a4 was in the S5 and S6 helices in the transmembrane region (∼1.7Å root mean square deviation, Figure 8A-A’ and Figure Supplement 11A-C), which is reported to contain the ligand binding pockets of Ors (Del Mármol et al. 2021; Wang et al. 2024; Zhao et al. 2024). We then substituted each of the 32 amino acids in *S. flava Or42a4* individually with the corresponding residues from *S. flava* Or42a3 *in silico,* predicted the 3D structures of the chimeras, and aligned them with *S. flava* Or42a4 in 3D space until the local structural differences were resolved. Remarkably, the substitutions of A181D and S307P in *S. flava* Or42a3 (hereafter referred to as A181D S307P) reduced the root mean square deviation to approximately 0.1 Å in the S5 and S6 helices when aligned with *S. flava* Or42a4, indicating that these two mutations are likely to explain the local structural differences (Figure 8A-A’ and Figure Supplement 11A-C).

**Figure 8:**
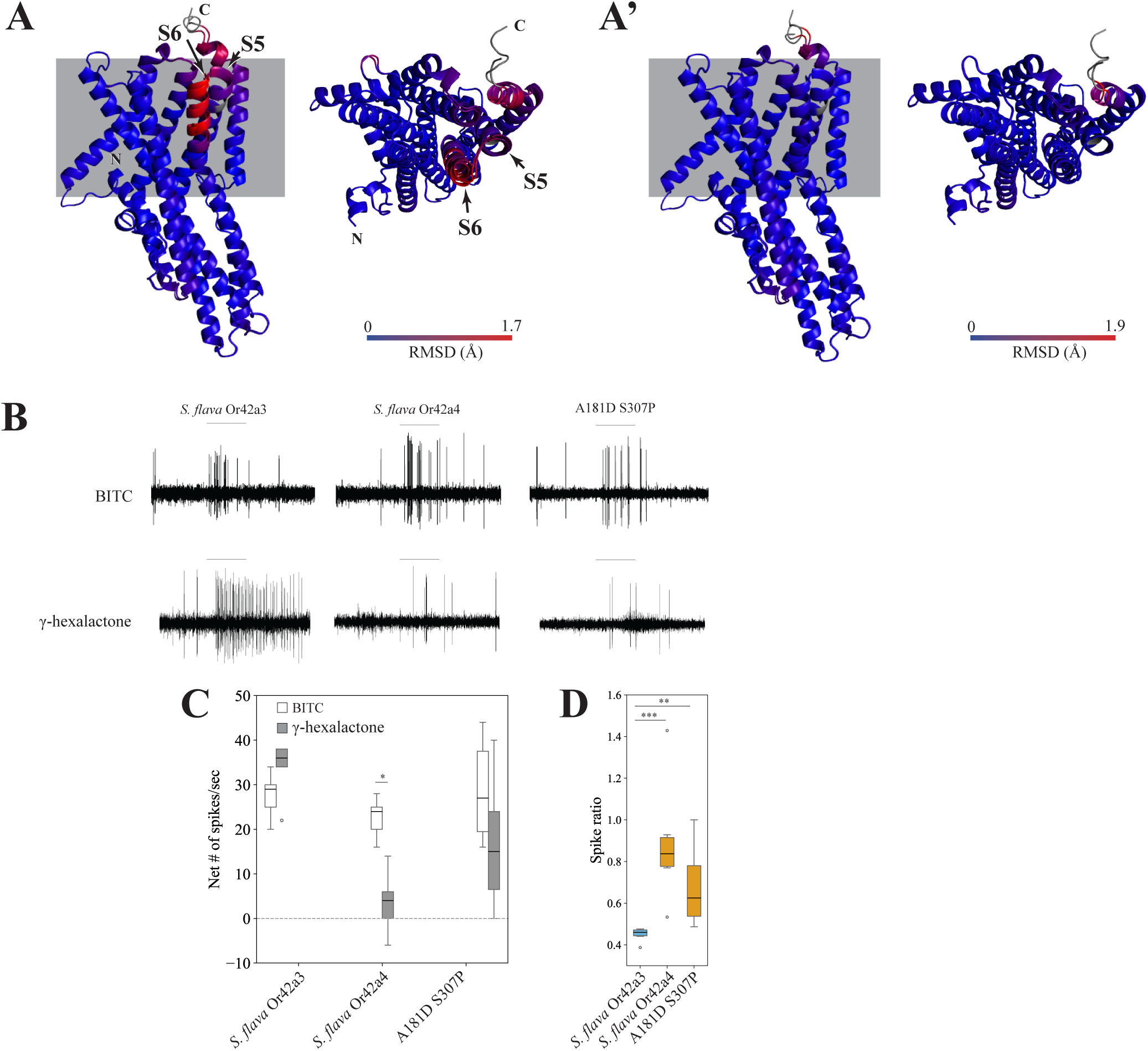
Two amino acids are critical for changing the sensitivity of *S. flava* paralogs from fruit volatiles to isothiocyanates. (A-A’) The 3D alignment of *S. flava* Or42a3 and *S. flava* Or42a4 predicted by AlphaFold2 **(A)**, and 3D alignment of *S. flava* Or42a4 and a chimeric Or42a with two amino acid substitutions (A181D and S301P) in the background of *S. flava* Or42a3 **(A’)**. The root mean square deviation is visualized with a color gradient from blue (low) to red (high) in both the side (left) and the top view (right). The upper and lower sections of the side view respectively represent the extracellular and the intracellular regions separated by the cell membrane (gray rectangles). **(B-D)** Representative single sensillum recording from *D. melanogaster* at1 sensilla expressing heterologous *S. flava* Or42a3, Or42a4, and the chimera (genotype: *UAS-Or42a/CyO; Or67d ^Gal4^*) upon stimulation with butyl isothiocyanate (BITC) and γ-hexalactone (**B**), at1 population responses to BITC (white bars) and to γ-hexalactone (gray bars) (**C**, n=6-10 sensilla from 3-4 animals/genotype; * p<0.05, Mann-Whitney U tests), and spike ratio [**D**, (response to BITC) / (response to BITC + response to γ-hexalactone); **p<0.01, ***p<0.001, Kruskal-Wallis ANOVA followed by Dunn’s multiple comparisons]. The odor tuning of the chimera, with just two amino acid substitutions, recapitulates that of *S. flava* Or42a3.

We then used the *D. melanogaster* at1 empty neuron system to investigate whether these two amino acid substitutions could account for the differences in odor sensitivity between *S. flava* Or42a4 and *S. flava* Or42a3. The A181DS307P variant and the two *S. flava* paralogs showed similar moderate responses to butyl ITC (Figure 8B-C). However, the butyl ITC to γ-hexalactone response ratios of *S. flava* Or42a4 and A181D S307P were not different from each other, but were both higher than the response ratio of *S. flava* Or42a3 (Figure 8D). These findings suggest that the two amino acid substitutions in *S. flava* Or42a4 are sufficient to increase the sensitivity to butyl ITC relative to γ-hexalactone. This effect was observed in flies carrying the heterozygous genotype A181D S307P/+ but not in flies with the homozygous A181D S307P genotype (Compare Figure 8C-D with Figure Supplement 12), possibly due to response saturation or dominant-negative effects. In summary, our findings demonstrate that the A181D and S307P substitutions in *S. flava* Or42a4 are critical for shifting the receptor’s sensitivity from fruit odorants to ITCs.

## 3. DISCUSSION

In this study we addressed three major questions: (1) Do non-specialist insects use olfaction to detect and avoid toxic isothiocyanate (ITC) compounds released by mustard plants? (2) Was this ancestral olfactory capability co-opted and expanded in mustard plant specialists for facilitating hostplant detection? (3) What molecular changes underlie the olfactory receptor sensitivity shift from fruit odors to hostplant-derived toxins? Answering these questions is important for understanding which molecular changes in the chemoreceptors of ancestral non-specialists enabled adaptation to toxic ecological niches in derived species, a central issue in the evolution of herbivory. We found that plant-derived ITCs are detrimental to microbe-feeding *D. melanogaster* through volatile exposure, and that the olfactory receptor Or42a is necessary for its detection and for behavioral aversion. To our knowledge, this is the first ITC-olfactory detector in *D. melanogaster*. In the mustard plant specialist *S. flava*, homologous olfactory receptors are triplicated and have an expanded ITC-sensitivity range, accompanied by an increase in the number of ITC-detecting OSNs. Finally, we discovered that two amino acid changes are sufficient to shift the odor sensitivity of these paralogs from fruit odors to ITC volatiles.

### 3-1. The plant-derived volatile allyl isothiocyanate is toxic to *D. melanogaster* and its detection and avoidance is mediated by *Or42a*-positive olfactory sensory neurons

Plants have evolved a diverse array of specialized metabolites, including electrophilic ITCs, that repel or intoxicate insects (Ibanez et al. 2012; Noge and Becerra 2015; Tocmo et al. 2021). ITCs are not only detected by contact (Bell et al. 2018), but some mustard-plant specialists such as the diamondback moth *Plutella xylostella* (Liu et al. 2020) and *S. flava* (Matsunaga et al. 2021) also detect these compounds via olfaction. However, it was unclear whether non-specialists such as *D. melanogaster* have evolved strategies to detect these toxic compounds using olfaction. We found that the olfactory receptor Or42a is necessary for OSN responses in the sensilla where it is expressed (palp basiconic 1a sensilla, Figure 3C). Furthermore, this Or was necessary for inducing olfactory aversion to AITC in two different behavioral contexts (Figure 4 and Figure Supplement 2A). Thus, in *D. melanogaster*, Or42a works in combination with the “wasabi” taste receptor” TrpA1 and Painless (Al-Anzi et al. 2006; Kang et al. 2010; Mandel et al. 2018), and possibly with other Ors (see next paragraph), to facilitate adaptive behavioral avoidance of these chemicals. It remains to be tested whether other non-specialist organisms across phyla also possess olfactory sensors tuned to volatile isothiocyanates.

We found that Or42a is also necessary for behavioral attraction to the fruit-derived volatile γ-hexalactone (Figure Supplement 2D-E and J). How does a single olfactory channel mediate aversion to AITC while also driving attraction to γ-hexalactone? Our experiments suggest that simultaneous activation of Or42a and the “generalist” OR7a (which is housed in OSNs in ab4 and respond to aversive odorants, Lin et al., 2015) could mediate aversion to AITC (Figure 4 and Figure Supplement 2F-G, M). Similarly, simultaneous activation of Or42a and Or35a (which is housed in ac3b, responds to γ-hexalactone and mediates attraction to yeast odors, Yao et al., 2005; Oh et al., 2021) could mediate attraction to γ-hexalactone (Figure Supplement 2H-I, M). Future investigations at the circuit level, particularly on the role of interglomerular interactions via local neurons in the insect primary olfactory center (Haverkamp et al. 2018), should further elucidate the neural mechanisms mediating these behavioral responses of opposite valence.

### 3-2. Duplication and functional evolution of the olfactory receptors *Or42a* and *Or67b* in *Scaptomyza* mustard plant specialists

Brassicales specialist need to detect a wide range of ITC compounds for effective host plant location, as these plants release species-specific volatile ITCs at particular ratios and concentrations (Wu et al. 2021). We previously reported that *S. flava* also has triplicated and positively selected *Or67b* copies (Matsunaga et al. 2021). The Ors encoded by these paralogous *Or67b* copies respond to aromatic and some aliphatic ITCs in a paralog-specific manner, while the *D. melanogaster* and *S. pallida* Or67b single copies did not respond to any volatile ITC compound (Matsunaga et al. 2021). However, all three *S. flava* Or67b paralogs showed poor responses to organosulfur ITCs, including AITC. In this study, we found that *S. flava* also expresses tandem triplicates of the *Or42a*s (*Or42a2*, *Or42a3*, and *Or42a4*; Figure 6A and Figure Supplement 6). Furthermore, OSNs housed in the pb1a-like sensilla of the mustard specialist species responded to many volatile ITCs, including AITC (Figure 5 and Figure Supplement 5). Thus, the gene duplications and amino acid substitutions of Or42a, along with those of Or67b, both likely play an important role in enabling Brassicales-plant specialist *Scaptomyza* species to detect a wide range of ITCs. Furthermore, mustard plant specialization is likely aided by the losses of genes encoding four olfactory receptors that detect fermentation odors in *D. melanogaster* and are necessary for attraction to these odors, and by the loss of an ancestral olfactory receptor (*Or7a*, housed in ab4a) that mediates aversion to AITC (Figure Supplement 2F-G; Goldman-Huertas et al. 2015).

What is the functional relevance of ITC-sensitive Ors expressed in two different olfactory organs? Or67b in *S. flava* is primarily expressed in the antennae (Matsunaga et al. 2021), whereas Or42a in *S. flava* and the other drosophilids is expressed in the maxillary palps (Figure 6B). In *D. melanogaster*, maxillary palps OSNs have lower sensitivity thresholds to certain host-related compounds compared to antennal OSNs (Dweck et al. 2016). Indeed, *S. flava* Or42a paralogs are much more sensitive to AITC than the Or67b paralogs (compare Figure 5 with Figure 4 in Matsunaga et al. 2021). Odor response redundancy between antennal and maxillary palp Ors could have evolved to further underpin olfactory orientation over both long and short distances in drosophilids (Dweck et al. 2016). Given the proximity of Or42a OSNs to the mouthparts, their activation could potentially modulate feeding behaviors, as suggested in *D. melanogaster* (Shiraiwa 2008). *S. flava* and *S. montana* females feed on the juice that seeps into the leaf wounds they create in Brassicales plants before oviposition (Peláez et al. 2022). Given this stereotyped feeding behavior, while the activation of contact chemoreceptors by ITCs could help females assess the suitability of an oviposition site through taste, flies may also be aided by maxillary palp olfactory activation even before tasting the plants.

Although the OSNs in pb1a sensilla of the two mustard specialist species have a broad ITC response range (Figure 5), *S. flava* has triplicated *Or42a*s but *S. montana* has only one copy (Figure Supplement 6). This suggests that both the mustard plant specialization and the mutations underlying the expanded ITC sensitivity range of Or42a preceded the triplication of *Or42a*. What is then the adaptive value of the *Or42a* triplication? We observed that *S. flava* pb1a OSNs, which express at least one of the three triplicated Or42as (Figure 6A-B), exhibited reduced sensitivity to fruit-borne γ-hexalactone in comparison with *S. montana* pb1a OSNs (Figure Supplement 5). Only *S. flava* Or42a4 acquired the two key mutations that increased the ITC to γ-hexalactone response ratio, as evidenced by the weaker γ-hexalactone response of the chimeric Or compared to that of *S. flava* Or42a3 (Figure 8). In agreement with these observations, structural alignment of the 3D models predicted by AlphaFold2 revealed that *S. montana* Or42a2 aligned well with *S. flava* Or42a3, but not with *S. flava* Or42a4 (Figure Supplement 11D). Based on these findings, we hypothesized the following sequence in the evolution of Or42a (Figure Supplement 13): (1) mustard specialization and broadening of ITC sensitivity in the ancestral drosophilid Or42a that was already sensitive to some ITCs like AITC; (2) speciation and triplication of Or42a in *S. flava* but not in *S. montana*; (3) relaxation of evolutionary constraints due to gene duplications, allowing *S. flava* Or42a4 to reduce its sensitivity to γ-hexalactone. It was suggested that *S. flava* has a broader host range (Maca 1972, Martin 2004) than *S. montana*, which requires indolic glucosinolates to use plants as hosts (Gloss et al. 2017). Thus, although mustard specialization preceded the Or42a duplication, we hypothesize that gene duplication was an important event in driving adaptation to new ecological niches in herbivorous *Scaptomyza*.

### 3-3. Evolution of specialized olfactory receptors is coupled with expansion of maxillary palp sensilla and its associated olfactory sensory neurons

We found that OSNs housed in pb1a sensilla from the mustard plant specialists *S. montana* and *S. flava* respond to a broad range of ITC compounds (Figure 5 and Figure Supplement 5), and that this sensilla type was numerically expanded in these two species (Figure Supplement 6-7). Concomitantly with this, we discovered an increase in the number of *Or42a*-positive OSNs in *S. flava* compared to *S. pallida* (Figure 6). Similar increases in OSNs which detect odors that bear species-specific biological significance have been reported in several *Drosophila* species. For instance, the noni fruit specialist *D. sechellia* and the seasonal specialist of “screw pine” (*Pandanus* spp.) fruits *D. erecta* both show an increase in the number of *Or22a-*positive OSNs (Dekker et al. 2006; Linz et al. 2013; Auer et al. 2020). In *D. sechellia*, these OSNs enhance odor tracking by reducing adaptation in second-order projection neurons (Takagi et al. 2024). We hypothesize that the increase in the number of *Or42a*-positive OSNs in *S. flava* may similarly contribute to enhance odor sensitivity and tracking during host plant finding, although this remains to be investigated.

### 3-4. Insight into the binding pocket of *S. flava* olfactory receptor Or42a paralogs

Our results demonstrate that the Or42a paralogs from microbe-feeding species show strong responses to both AITC and γ-hexalactone (Figure 7), while the paralogs from the herbivorous *S. flava* show a notable shift in olfactory sensitivity, which aligns with the fact that this species has undergone a full transition to herbivory. For example, *S. flava* Or42a3 showed a moderate response to γ-hexalactone (although about an order of magnitude lower than that of the Or42a from the microbe-feeding *S. pallida*), and *S. flava* Or42a4 was even less sensitive (Figure 7). Furthermore, *S. flava* Or42a4 had a relatively higher sensitivity to butyl ITC compared to Or42a3 (Figure 7). To explore the mechanisms underlying this shift from fruit-detector to ITC-detector, we employed computational and functional approaches to identify the amino acid substitutions responsible for the differencial odor sensitivity of *S. flava* Or42a3 and Or42a4.

Our AlphaFold2-led screening, combined with site-directed mutagenesis and electrophysiological studies, identified two key substitutions in the transmembrane region of Or42a4, A181D and S307P, that are critical for the odor sensitivity switch (Figure 8). Proline is a secondary structure breaker (Chou and Fasman 1978; Levitt 1978), and the substitution of serine by proline (S307P) likely influenced the conformation change of the binding pocket, altering the protein structure, polarity, and hydrophobicity. Similarly, the substitution of alanine by aspartic acid (A181D) can alter protein polarity and hydrophobicity. Thus, these two amino acid changes likely account for the notable change in ligand sensitivity. However, this change in ligand sensitivity was observed only in flies heterozygous for A181D and S307P (Or67d^Gal4^; UAS-A181D S307P/+), but not in homozygous flies (compare Figure 8C-D with Figure Supplement 12). Given that homozygous flies express A181D and S307P more strongly than the heterozygous flies, it is possible that the OSNs’ response to both odorants reached saturation in the former, obscuring differences in spike ratios. Alternatively, excessive expression of A181D and S307P may have altered the 3D structure of the heteromeric protein, possibly through dominant-negative effects between the copies, reverting it closer to the original *S. flava* Or42a3 conformation and restoring the original binding pocket. Nonetheless, the partial rescue we observed in heterozygous flies indicates that A181D and S307P are critical for altering the binding pocket structure, enabling the Or to better accommodate butyl ITC instead of γ-hexalactone. Although the remaining amino acid substitutions in Or42a4 might alter e.g. signal transduction and/or receptor stability, it is remarkable that only two substitutions out of 32 amino acid differences between paralogs were sufficient to shift ligand sensitivity.

## 4. CONCLUSIONS

Taken together, our findings reveal that non-herbivorous, microbe-feeding insects like *D. melanogaster* have evolved promiscuous olfactory sensory mechanisms that allow them to detect and avoid specialized plant-derived volatile electrophilic toxins, such as isothiocyanates. In contrast, Brassicales plant specialists like *S. flava* not only have physiological adaptations to detoxify these toxic compounds but have undergone significant evolutionary sensory adaptations for aiding host plant location, including turning shifts and expansions of specialized isothiocyanate olfactory receptors. Furthermore, our use of AlphaFold2, followed by site-directed mutagenesis and electrophysiology, identified critical amino acid changes for the evolution of these specialized odorant receptors that we confirmed experimentally. Thus, ancestral Ors that mediate toxin aversion in generalist species can be co-opted and diversified in derived specialists through gene duplications and tuning shifts facilitated by relatively few amino acids substitutions.

## 5. MATERIAL AND METHODS

### Fly Husbandry

*D. melanogaster* was reared on cornmeal medium. Microbe-feeding *S. pallida* (*S. pallida* subgenus *Parascaptomyza*) and *S. hsui* (*S. hsui*subgenus *Hemiscaptomyza*) were reared in cornmeal molasses media covered with a mixture of Carolina biological supply instant *Drosophila* media (Burlington, NC) mixed with blended spinach leaves, and then covered with a layer of defrosted frozen spinach leaves. The obligate leaf-miners *S. flava* and *S. montana* (subgenus *Scaptomyza*) were cultivated on potted fresh laboratory-grown *Arabidopsis thaliana* Col-0. Isofemale lines of microbe feeding *S. hsui* and *S. pallida*, as well as the herbivorous *S. montana,* were collected along Strawberry Creek on the UC-Berkeley Campus in Berkeley, California, USA (Kim et al. 2021), and a line of *S. flava* was collected from a meadow near Dover, New Hampshire, USA. All species were kept at 23±2 ℃ and 60% relative humidity in a 10:14 hours light-dark cycle under fluorescent lights. The following lines (stock #) were obtained from the Bloomington *Drosophila* Stock Center: *Or42a-/-* (60821), *Or42a-Gal4* (9970), *w^1118^* (3605), *TrpA1^1^* (26504), *Or7a-/-* (91811), and a genetic background control for the three *Or* null mutant lines (68384). The *Or35a-/-* line (10564) was obtained from the Korea Stock Center. The *Or67d^Gal4^* line was a gift from the laboratory of Barry J. Dickson.

### Single sensillum recordings (SSRs)

One to five days old fed female flies were prepared for SSR as described in Matsunaga et al. (2021). Briefly, a silicon tube delivering a constant flow of charcoal-filtered air (16 ml/min, measured using a flowmeter, Gilmont Instruments, USA) was placed near the fly’s head capsule, and the tip of the stimulation pipette (50 ml) was inserted into the constant air stream. The stimulation pipette contained a 0.5 cm x 5 cm piece of filter paper loaded with 20 µl of an odorant solution or the solvent control. A pulse of clean air (duration = 1 sec) was delivered to the stimulus pipette using a membrane pump operated by a Stimulus Controller CS 55 (Syntech, Buchenbach, Germany). Sensilla identification was conducted using the following diagnostic odorants (as described in Dweck et al. 2016; Gonzalez et al. 2016), all >95% pure (Sigma-Aldrich, St. Louis, MO, USA): ethyl acetate (CAS# 141-78-6) for identifying *D. melanogaster* pb1a; AITC (CAS # 57-06-7) for identifying *Scaptomyza* pb1a-like OSNs; 4-ethylguaiacol (CAS# 2785-89-9) for identifying *D. melanogaster* and *Scaptomyza* pb1b and pb1-like OSNs; fenchone (CAS# 1195-79-5) for identifying *D. melanogaster* and *Scaptomyza* pb2a and pb2a-like OSNs; guaiacol (CAS# 90-05-1) for identifying *D. melanogaster* and *Scaptomyza* pb2b and pb2b-like OSNs; phenethyl acetate (CAS# 103-45-7) for identifying *D. melanogaster* pb3b, *S. flava* pb3b-like, and *S. montana* pb3b-like OSNs; and 2-heptanone for identifying *S. hsui* and *S. pallida* pb3a-like OSNs. All odorants were diluted in mineral oil (CAS# 8042-47-5) except γ-hexalactone (CAS # 695-06-7), which was diluted in dimethyl sulfoxide (CAS# 67-68-5) because it did not dissolve completely in mineral oil and sometimes produced response artifacts. Odorants were diluted to 1:100 vol/vol for stimulation unless otherwise noted. Supplementary file 1 lists all the chemicals used in this study.

The “net number of spikes/second” was obtained by counting the number of spikes originating from the OSN of interest within a 0.5-second timeframe which started 0.2 seconds after the onset of stimulation. This count was then adjusted by subtracting the background spiking activity (# of spikes within a 0.5 second interval preceding the onset of the stimulation) and then doubled to represent the number of spikes/second. In all figures, unless otherwise stated, we represent the “control-subtracted net # of spikes/sec” to odorant stimulation, calculated by subtracting the average net # of spikes/sec in response to the solvent control (mineral oil or dimethyl sulfoxide) from the net # of spikes/sec evoked by each odorant stimulation. Control-subtracted spike data are compiled in Supplementary file 2. The butyl ITC to γ-hexalactone spike ratio (Figure 8D and Figure Supplement 12) was calculated as: net # of spikes/sec evoked by butyl ITC / (net # of spikes/sec evoked by butyl ITC + net # of spikes/sec evoked by γ-hexalactone). We used this denominator for the ratio because the control-subtracted net # of spikes upon γ-hexalactone stimulation occasionally produced negative values (likely a response to the solvent control).

Half maximal effective concentrations (EC_50_) were calculated using *Quest Graph™ EC50 Calculator* (AAT Bioquest, Inc., 4 Mar. 2025, https://www.aatbio.com/tools/ec50-calculator). We primarily used the two-parameter feature with normalization, where responses were normalized to the largest response within the same chemical-species pair, with the minimum set to 0. This approach was chosen because the lowest tested concentration (10^-5^) still elicited non-zero spike activity (>10) in some chemical-species pairs. The four-parameter method without normalization, in which the maximum and minimum responses were free parameters, was used in cases where the two-parameter method failed to fit a logistic regression (summarized in Supplementary file 2) or when analyzing *sec*-butyl ITC data. For this odorant, the four-parameter method was necessary because the highest tested concentration (10^-2^) did not reach saturation (<100 spikes). When both the two-parameter and four-parameter methods failed due to lack of convergence at the lowest concentration (10^-5^), resulting in a calculated value of 0, we instead used the minimum value observed within that chemical-species group.

The *Or67d*^GAL4^ line was used to generate flies expressing *Or42a* homologs in the at1 “empty neuron” system (Kurtovic et al. 2007). We selected at1 (instead of the more commonly used antennal basiconic ab3 “empty neuron system”, Dobritsa et al. 2003) because some insects use host-derived chemicals as pheromones (Reddy and Guerrero 2004), and in *D. melanogaster* pheromones sometimes activate OSNs housed in trichoid sensilla only (Xu et al., 2005; Benton et al., 2007).

The spike amplitude differences between the two types of OSNs housed in pb3a-like and pb3b-like sensilla were less distinct in *Scaptomyza*, and therefore we could not completely rule out the possibility that we occasionally erroneously assigned spiking to each of these two OSNs types.

### Immobility assay

To investigate the effect of AITC volatiles in wild-type *D. melanogaster* (Canton-S strain), we used a 9 cm diameter plastic petri dish (Nunc, Denmark) with a piece of fabric mesh placed horizontally between the base and the lid, creating two chambers (Figure Supplement 1A). The upper chamber housed 8-10 male flies 3-5 days old, and the lower chamber contained four 5 µl drops of the odor solution (or the solvent control) evenly dispersed. Because the mesh prevented the flies from reaching the bottom chamber, insects were exposed to the volatile chemicals but could not directly contact (i.e. taste) the odor solution, unless the molecules adhered to the walls of the chamber after volatilizing. After each test started, we counted the number of mobile flies every 10 minutes up to one hour and calculated the percentage of mobile flies at each time point. Flies exhibiting no movement for >30 seconds were likely intoxicated. AITC and γ-hexalactone were respectively diluted in either mineral oil or dimethyl sulfoxide at various concentrations. Mobility data analysis was performed using the log-rank (Mantel-Cox) test (Mantel 1966). The complete immobility assay dataset is compiled in Supplementary file 3.

### Positional olfactory assay

To study the olfactory orientation of insects towards odors, we conducted assays with non-starved 3-4 days old mated females (Figure Supplement 1B). Flies (n=10-12/test) were anesthetized on ice (5-7 minutes) and placed in a small piece of clear Tygon tube, capped in both sides with a conical PCR plastic Eppendorf. After another about 4-5 minutes, the Tygon tube with the anesthetized flies was uncapped and connected to the cut ends of two glass Pasteur pipettes and the assay started; flies usually resumed activity after about 3-4 minutes. Each of the two opposite ends of the pipettes were respectively connected to a 1.75-ml glass vial containing 10 µl of the odor solution (AITC 1: 500 vol/vol, or γ-hexalactone 1:100 or 1:10 vol/vol) or 10 µl of the control solvent (mineral oil or dimethyl sulfoxide). Tests with apple cider vinegar used 30 µl instead, and water was used as a control. The distal ends of the pipettes were separated from the glass vials with a small piece of fabric mesh, which allowed the odorant to diffuse into the pipettes while also preventing insects from contacting the odor source (Figure Supplement 1B). The odor and control sides were switched between assays. Assays were conducted on a white surface under white light at 21-24 ℃, about 2-6 hours after lights on. Once each assay started, the number of flies in the pipette closest to the vial with the odor (referred to as “odor side” or “odorous tube” for simplicity), in the pipette closest to the vial with the solvent control (“control” side/tube), and in the Tygon tube that connected both pipettes (release site) were counted every 5 minutes until minute 35, and then again at 65 minutes in the case of tests with AITC. For each assay, we calculated the % of insects that made a choice for one or the other tube as: [(#of insects in the odor side + # of insects in the control side) / total number of insects released] x 100. The percentage of insects that choose the odorous tube was calculated based on the total number of insects that made a choice as: [# of insects in the odor side / (# of insects in the odor side + # of insects in the control side)] x 100. Assays in which less than 40% of insects made a choice for either side at all time points were discarded (<5% of assays). For each fly genotype and odorant, the % of insects that choose the odor side at each time point was compared against the median value expected under the null hypothesis that insects distributed at random between the two tubes (50% of insects in each tube) using one-sample signed rank tests. Thus, we assessed whether the insects significantly avoided the odorous tube (if median<50%), preferred it (if median>50%), or showed a random preference (median ca. 50%). In some cases, at each time point and for each odor (and concentration when applicable), the responses of the null mutants (*Or42a-/-*, *Or7a-/-* and *Or35a-/-*) and their genetic background control (line 6834 listed above) tested in parallel (i.e. in the same days) were compared via Mann-Whitney U tests. In all cases results were considered significant if p<0.05. In most cases we used two-tailed tests (e.g. for testing median_1_ ≠ median_2_), but in a few cases we used one-tailed tests (e.g. for specifically testing whether median_1_ > median_2_ or whether median_1_<median_2_, Figure Supplement 2F-I). The positional olfactory assay data is included in Supplementary file 4. In this and all behavioral assays, we used the line # 6834 instead of the more standard *w1118*, because the latest have visual defects that interfere with normal behavior (Ferreiro et al. 2017).

### Consumption assay

This behavioral assay (described in Reisenman and Scott 2019) measures if the presence of an odorant affected consumption of an appetitive solution. Groups of 2-4 days old mated female flies (n=11-15) were wet-starved for 24 hours and then transferred to a vial containing a piece of filter paper (2.7 cm diameter, Whatman, cat. No 1001 125) impregnated with 160 µl of 50 mM D-glucose (Sigma-Aldrich, USA) dyed blue with Erioglaucine (0.25 mg/ml, Sigma-Aldrich, St. Louis, MO, USA; Figure Supplement 1C). Flies were allowed to feed for 15 minutes (10 minutes in tests with γ-hexalactone), frozen (>60 minutes), and the amount of blue dye in the flies’ abdomen was scored blind to treatment (see below). The odor source consisted of a strip of filter paper (0.25 cm wide x 1.5 cm long) impregnated with either 10 µl of an odorant solution (test) or 10 µl of the solvent (control), which was placed inside a container (1.3 cm long x 0.75 cm diameter) with a meshed bottom affixed to the vial’s flug (Figure Supplement 1C). This allowed diffusion of odors into the fly vial but prevented flies from contacting the odor source. Control tests, with vials containing food solution but only the solvent control inside the meshed container, were conducted in parallel with experimental tests to control for fly cohort and day-to-day variability.

Food consumption was estimated by scoring individual flies in each vial blind to treatment using the following five-point scale (Reisenman and Scott 2019): 0 (no dye = no food), 0.25 (“trace” of blue dye), 0.5 (up to ¼ of the abdomen dyed blue), 1 (more than ¼ but less than ½ of the abdomen dyed blue), and 2 (more than ½ of the abdomen dyed blue). For each vial, a single feeding score value was calculated as: [(0 x n_0_ + 0.25 x n_0.25_ + 0.5 x n_0.5_ + 1 x n_1_ + 2 x n_2)_ / N], where n_(0-2)_ denotes the number of flies in each score category, and N the total number of flies/vial. Feeding scores from each test vial (flies offered food in presence of an odor) were normalized to the averaged feeding score of control vials (flies of the same genotype offered food in absence of odor) assayed on the same day. Normalized feeding scores for each genotype and odor were compared against the null hypothesis (median feeding score = 1) using one sample signed rank tests. That is, medians not different from 1 indicate that the odorant did not reduce neither enhanced consumption, while medians significantly less than 1 or more than 1 respectively indicate feeding aversion and enhancement. Normalized data from control and mutant flies were compared using Mann-Whitney U tests. In all cases results were considered statistically significant if p<0.05. The consumption assay data is compiled in Supplementary file 5.

### RNA-sequencing of maxillary palps

Newly emerged adults of *S. flava* and *S. pallida* were collected from our colony and kept in humidified vials with 10% honey water until dissection, to minimize potential differences in nutrition resulting from differences in the two species’ larval diet. Three-ten days old flies were anesthetized with CO_2_, and their maxillary palps were hand-dissected using forceps. Approximately 100-120 flies were pooled for a single sample. The dissected tissues were directly collected in LB+TGA lysis buffer from Reliaprep RNA Tissue Miniprep System (Promega, USA), and homogenized using a Biomasher Standard homogenizer (Takara Bio Inc., USA) in a dry ice ethanol bath. The sample lysates were stored at -80℃ until RNA extraction. Total RNAs were extracted from the lysates using ReliaPrep RNA Tissue Miniprep System (Promega, USA) according to the manufacturer’s protocol, and quantified using a Qubit RNA High Sensitivity kit (Thermo Fisher Scientific, USA). Library preparation was performed at the Functional Genomics Laboratory at UC Berkeley. Due to the low yields of our maxillary palp-derived total RNAs, cDNA libraries were first produced by Takara SMART-Seq mRNA Ultra-low input RNA kit (Takara Bio Inc., USA) with 8 cycles of PCR for the amplification, and then processed by KAPA HyperPrep kit for DNA (Roche Sequencing, USA) with 9 cycles of PCR for attaching in-house sequencing adapters and index primers. cDNA libraries were then sequenced on an Illumina NovaSeq 6000 150 PE S4 flowcell, targeting 25M read pairs per library by the UC Berkeley Vincent J. Coates Genomics Sequencing Laboratory. For read mapping, we used previously reported reference genome assemblies and gene annotations from *S. flava* (Peláez et al. 2023) and *S. pallida* (Kim et al. 2021) for subsequent bioinformatic analyses. Raw RNA-seq reads were filtered using Fastp v0.21.0 (Chen et al. 2018) and mapped to the respective reference genomes using STAR v2.7.1a (Dobin et al. 2013) to generate multiple alignment (BAM) files, which were then converted to read count data using HTseq v0.9.1 (Anders et al. 2015). Count data for the *Or* gene family were converted to RPM (reads per million; Supplementary file 6).

### Hybridization chain reaction RNA Fluorescent in situ hybridization (FISH)

One-four days old female *S. flava* and *S. pallida* were collected and anesthetized with CO_2_. Whole mouthparts were removed and immediately placed in 2 mL of fixative (4% vol/vol paraformaldehyde in 1x phosphate buffer saline with 3% vol/vol Triton X-100 added, PBST) in LoBind Eppendorf tubes, and fixed for 22 hours at 4°C on a nutator. For HCR RNA fluorescent in situ hybridization, we followed the manufacturer’s instruction (Molecular Instruments, Inc., Los Angeles, CA).

Samples were stained with 300 nM DAPI in 0.1% PBST for 15 minutes and then washed 3 times with 0.1% PBST for 5 minutes. Tissues were transferred to a microscope slide and mounted in a drop of ProLong diamond antifade mounting (Life Technologies Corp., Eugene, OR) and stored at 4°C until examination. Confocal imaging of fixed samples was performed using a Zeiss LSM 880 microscope in the AiryScan mode. Raw images were processed using Zeiss ZEN Black software. Orco-positive cells (visualized with the 488 nm laser) and Or42a-positive cells (visualized with the 633 nm laser) were manually counted using the “Cell Counter” plugin in Fiji (ImageJ) software. Supplementary file 7 contains cell counts from RNA Fluorescent in situ hybridization experiments.

### *Scaptomyza Or42a* gene cloning and generation of *UAS* lines

RNA was isolated from 15-25 days old laboratory-reared adults of both sexes of *S. pallida* and *S. flava* using the ReliaPrep Miniprep system (Promega, W.I., USA). Complementary DNA (cDNA) was synthesized using the qScript cDNA Supermix (Quantabio, Beverly, MA, USA). The paralogs were amplified via touchdown PCR, using primers that target the highly variable regions of the 5’ and 3’ untranslated regions (UTRs). The single *Or42a2* gene of *S. pallida* was amplified using touchdown PCR with primers targeting the coding region (Q5 DNA Polymerase, #M0491, NEB) (Supplementary file 8). All PCR products underwent gel purification (11-300C, Zymo Research) and were subsequently cloned using the Gibson Assembly (E2611S, NEB) following the manufacturer’s instructions into the UAST-attB vector (DGRC Stock 1419; RRID:DGRC1419, Bischof et al. 2007). The ligation products were transformed into DH5α competent cells (T3007, Zymo Research). After confirming the sequences using Sanger sequencing, the 5’ and 3’ UTRs of the plasmids containing the *S. flava Or42a* paralogs were removed. This was achieved by first amplifying the coding region using PCR, followed by gel purification and ligation back into UAST-attB vector via Gibson Assembly. The resultant plasmids were verified using Sanger sequencing. Finally, these plasmids were microinjected into the attP40 site (P{nos-phiC31\int.NLS}X, P{CaryP}attP40, line # 25709, Rainbowgene) to create transgenic lines for *S. pallida Or42a*, *S. flava Or42a2*, *S. flava Or42a3* and *S. flava Or42a4*. All lines were confirmed by Sanger sequencing prior to experiments.

### Screening of candidate amino acids using AlphaFold2 3D structural prediction

CDS of *S. flava Or42a3* and *S. flava Or42a4* were confirmed by palp RNAseq data using IGV_2.16.0. CDSs of *S. flava Or42a3* and *S. flava Or42a4* were then used as inputs into ColabFold (Jumper et al. 2021; Mirdita et al. 2022). Then the output models ranked first (rank1) were selected, visualized, and 3D-aligned by using PyMol2.5.3. We focused on the regions where predicted local distance difference test scores exceed 70, as structures with lower values are often unreliable (Mariani et al. 2013). We focused on the S5-S6 transmembrane helices because previous studies suggested that the binding pockets of odorant receptors are located in the transmembrane region (Butterwick et al. 2018; Del Mármol et al. 2021; Zhao et al. 2024), and our amino acid alignment of *S. flava* Or42a3 and *S. flava* Or42a4 provided the highest root mean square deviation scores in this region. In silico substitutions of amino acids were performed individually on *S. flava* Or42a3, and the resulting sequences were then re-input into ColabFold, the model with rank1 was selected, and the structures were 3D-aligned with that of *S. flava* Or42a4. This process of in silico mutation and alignment was repeated until the root mean square deviation scores for the S5-S6 region were reduced to a value comparable to the other region (∼0.1 Å). All pdb files used in this study are included in Supplementary file 9.

### Site-directed mutagenesis

Conventional PCR was conducted using plasmids of *S. flava Or42a3* as backbone (Q5 DNA polymerase, #M0491, NEB; Supplemental File 1). Primers were designed to introduce the point mutations (Supplementary file 1). The PCR product underwent gel purification (11-300C, Zymo Research) and the methylated plasmids were digested with DpnI for 1 hour at 37℃ (QuickChange, Agilient Technology, USA). Ligation was performed by incubation at 16℃ for 30 minutes (DNA ligation kit Mighty mix, Takara, Japan) and the products were transformed into DH5α competent cells (Takara, Japan). After confirming the sequence using Sanger sequencing, the plasmid was microinjected into the attP40 site (P{nos-phiC31\int.NLS}X, P{CaryP}attP40, line # 25709, Rainbowgene) to create the mutant A181D S307P fly line. The mutations were confirmed by sequencing prior to experiments.

## ACKNOWLEDGEMENTS

We are grateful to two anonymous reviewers and to the editor for their constructive comments and suggestions that improved this manuscript. T.M. thanks Hany Dweck for valuable insights about conducting SSR of maxillary palp sensilla, to Dr. Akinao Nose and Dr. Hiroshi Kohsaka for their advice and encouragement, and to the members of the Nose’ laboratory and an anonymous researcher at the University of Tokyo for discussions. We thank members of the Whiteman and Scott Laboratories for discussions and comments on the manuscript. The Bloomington Drosophila Stock Center, the Korean Stock Center, and the Drosophila Genomics Resource Center (NIH Grant 2P40OD010949) kindly provided fly lines. We are grateful to T. Michael Keesey for the use of *A. thaliana* silhouette in PhyloPic. This work was supported by the Grant-in-Aid for Scientific Research Activity Start-up (T.M.), the Uehara Memorial Foundation (award number 201931028 to T.M.), the National Institute of Health (NIH) Grant (award number R35GM139653 to Gary Karpen), and the National Institute of General Medical Sciences of the NIH (award number R35GM119816 to N.K.W.).

## AUTHORS’ CONTRIBUTION

All authors contributed to the experimental design, analysis, and interpretation of results. T.M. initiated the project and performed all experiments, except for the behavioral experiments (Fig. 4, Fig. Supplement 2A-J, and others indicated in the text and/or Figure captions), which were conducted by C.E.R. RNA fluorescent in situ hybridization experiments (Fig. 6C-E) were performed by B.G.H. and S.R.; RNA-seq analysis (Fig. 6B) was performed by H.C.S. D.T. cloned *UAS-Or42a*s. J.W. contributed to the behavioral experiment presented in Fig. 4B. S.R.R. provided advice on SSR to T.M. T.M. wrote the manuscript in its original and revised versions with a substantial contribution from C.E.R.; C.E.R also wrote and revised the sections corresponding to the experiments she conducted. All authors, especially N.K.W. and C.E.R., provided feedback on the overall manuscript. N.K.W. supervised T.M. during the initial stages of the project.

## SUPPLEMENTARY MATERIAL AND METHODS

### Dilution series

*Trans*-2-hexenal, AITC, butyl ITC, isobutyl ITC, *sec*-butyl ITC, and ethyl butyrate were diluted in mineral oil, and γ-hexalactone was diluted in DMSO to concentrations ranging from 10^-5^ to 10^-2^ vol/vol. Half maximal effective concentration (EC50) was generated by GraphPad Prism v10.2.1.

### Construction of the Or42a gene tree

The amino acid sequences of *D. melanogaster Or42a,* and *S. flava Or42a1, S. flava Or42a2, S. flava Or42a3,* and *S. flava Or42a4* were obtained from Flybase (2024_01), and Goldman-Huertas et al. (2015). Blast searches were conducted to annotate *S. hsui Or42a1, S. pallida Or42a1, S. graminum Or42a1, S. graminum Or42a2, S. montana Or42a1, S. montana Or42a2, S. flava Or42a1, S. hsui Or42a2,* and *S. pallida Or42a2,* using *S. flava Or42a1* and *S. flava Or42a2* as query against the genomes of *S. hsui* (ASM1815282v1), *S. pallida* (ASM1815296v1), *S. graminum* (ASM1890183v1), and *S. montana* (ASM1890430v1). Amino acid sequences of *S. hsui Or42a1, S. pallida Or42a1, S. gramina Or42a1, S. graminum Or42a2, S. montana Or42a1, S. montana Or42a, S. flava Or42a1, S. flava Or42a2, S. flava Or42a3, S. flava Or42a4, S. hsui Or42a2, S. pallida Or42a2* (Katoh and Standley 2013), and *D. melanogaster Or42a* were aligned by MAFFT v7.511 with JTT as scoring matrix for amino acid sequences with L-INS-i strategy. Maximum-likelihood gene tree with 1000 bootstrap cycles using the above sequences were constructed with RAxML (v8.2.19) (Stamatakis 2014). GAMMA model of rate heterogeneity was applied. CDS and alignment of *Or42a* are included in Supplementary files 12 and 13, respectively.

### Alignment of *S. flava Or42a3* and *Or42a4,* and *S. montana Or42a2* using AlphaFold2 3D structural prediction

CDSs of *S. flava Or42a3*, *S. flava Or42a4,* and *S. montana Or42a2* were used as inputs into ColabFold (Jumper et al. 2021; Mirdita et al. 2022). The output models ranked first (rank1) were selected, visualized, and 3D-aligned by using PyMol2.5.3.

### Syntenic analysis using DiGAlign

To analyze synteny around the *Or42a2-Or42a4* loci, we extracted genomic regions spanning a total of 30kb (15kb upstream and 15kb downstream of the *Or42a2*s start site) from the genomes of *S. montana* (ASM1890430v1), *S. pallida* (ASM1815296v1), and *S. hsui* (ASM1815282v1). For *S. flava* (sfla_v2), we extracted a 60,199 bp region, encompassing 15kb upstream of the *Or42a2* start site through 15kb downstream of the *Or42a4* start site (Kim et al. 2021; Peláez et al. 2023). For visualization purposes, the genomes of *S. montana, S. graminum,* and *S. pallida* were reverse-aligned. Synteny maps and dot plots were generated and visualized using DiGAlign (Nishimura et al. 2024).

## SUPPLEMENTARY FILES

Supplementary file 1: Chemical list

Supplementary file 2: Control-subtracted net number of spikes/sec and EC_50_ obtained from SSRs

Supplementary file 3: Mobility assay data

Supplementary file 4: Olfactory positional assay data and trap assay data

Supplementary file 5: Consumption assay data

Supplementary file 6: Expression intensity from maxillary palp RNA-seq

Supplementary file 7: OSNs counts from RNA fluorescent in situ hybridization

Supplementary file 8: Primer list

Supplementary file 9: AlphaFold2 (Colabfold) prediction in pdb format

Supplementary file 10: Pb sensilla counts from single sensillum recording experiments

Supplementary file 11: CDS of *Or42a* homologs in fasta format

Supplementary file 12: Alignment of *Or42a* homologs in fasta format

## DATA AVAILABILITY

The data presented in this study are available on request from the corresponding authors.

## ABBREVIATIONS

AITC: allyl isothiocyanate
ITC: isothiocyanate
Or/OR: olfactory receptor
OSN: olfactory sensory neuron Pb: palp basiconic
SSR: single sensillum recording

---

**Figure Supplement 1:**
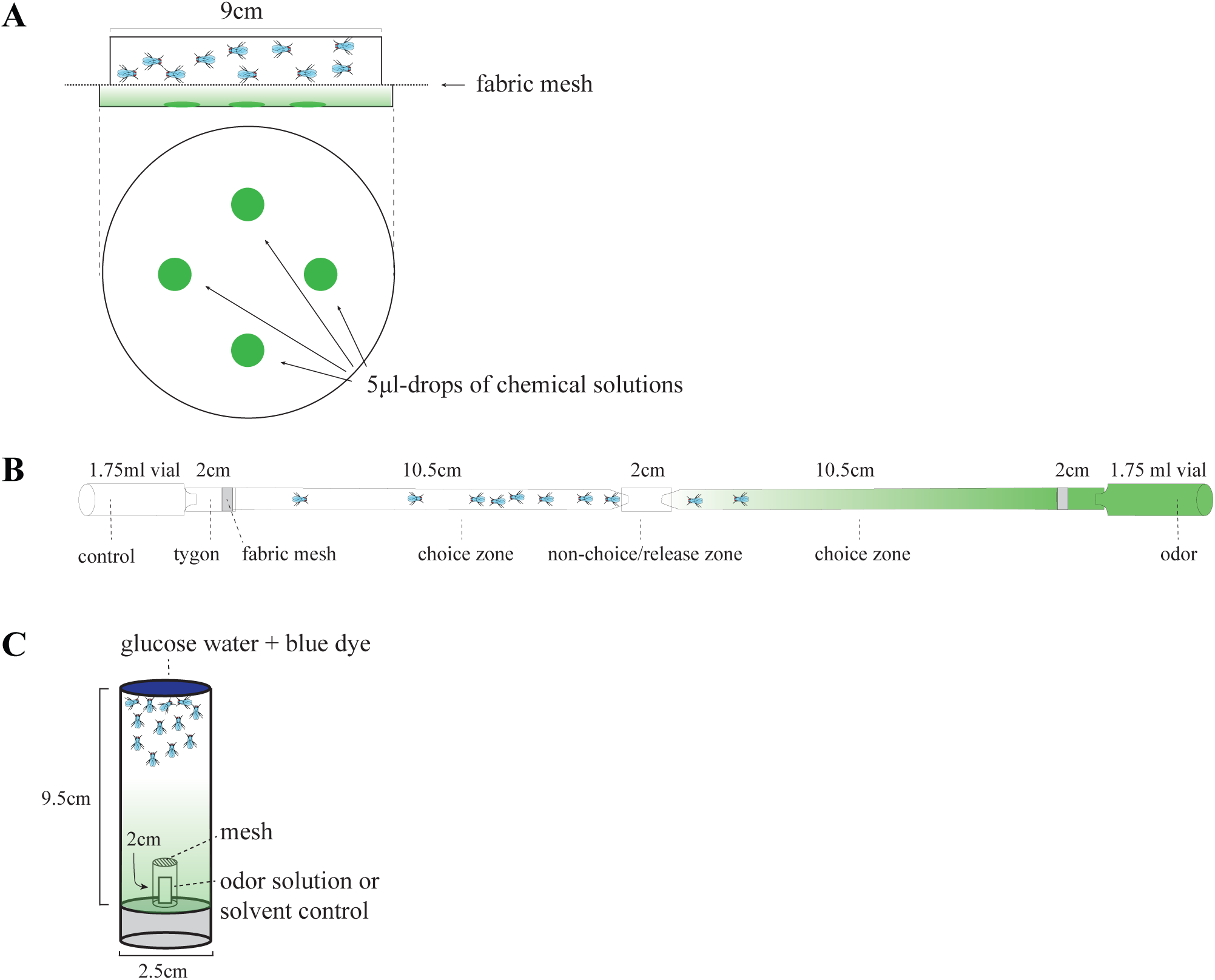
Schematic representations of the behavioral assays. **(A)** Mobility assay used as a proxy for intoxication. Mated female flies (n=10) were placed in the upper chamber (lateral view) and monitored every 10 minutes in presence of volatile chemicals (4 drops of a 5 µl solution in the lower chamber, top view). The top and the bottom chambers were separated by a fabric mesh, preventing direct contact between the flies and the chemical solutions. **(B)** Positional olfactory assay. Groups of 10-12 non-starved cold-anesthetized mated females were placed in a small piece of clear Tygon tube, which was then used to connect the narrow ends of two cut glass Pasteur pipettes (non-choice/release zone). The distal end of each glass pipette was connected to a glass vial containing 10 µL of the odor solution or the solvent control; a piece of fabric mesh prevented insects from entering the vials containing the odor or control solutions. The number of flies in each of the two tubes, as well as in the middle release section, were counted every five minutes until minute 35, and in tests with allyl ITC, again at 65 minutes. The % of insects that choose one or the other tube over the total number of insects released, and the % of insects in the tube closest to the odor source over the total number of insects that choose one or the other tube, was then calculated for each time point. **(C)** Feeding assay in presence of odors or the solvent control. In each test a group of 24 hours wet-starved mated female flies (n=11-15) was placed in a vial containing a piece of filter paper impregnated with 160 µl of 50 mM D-glucose dyed blue. A small container with a mesh bottom was affixed to the inner side of the vial cap contained a piece of filter paper loaded with 10 µl of an odorant solution or the solvent control, such as that flies could smell but not contact the odor source. After 15 minutes, vials were frozen for at least 60 minutes, the amount of blue dye in the abdomen of each fly in each vial was quantified blind to treatment, and a feeding score was calculated for each vial as explained in materials and methods.

**Figure Supplement 2:**
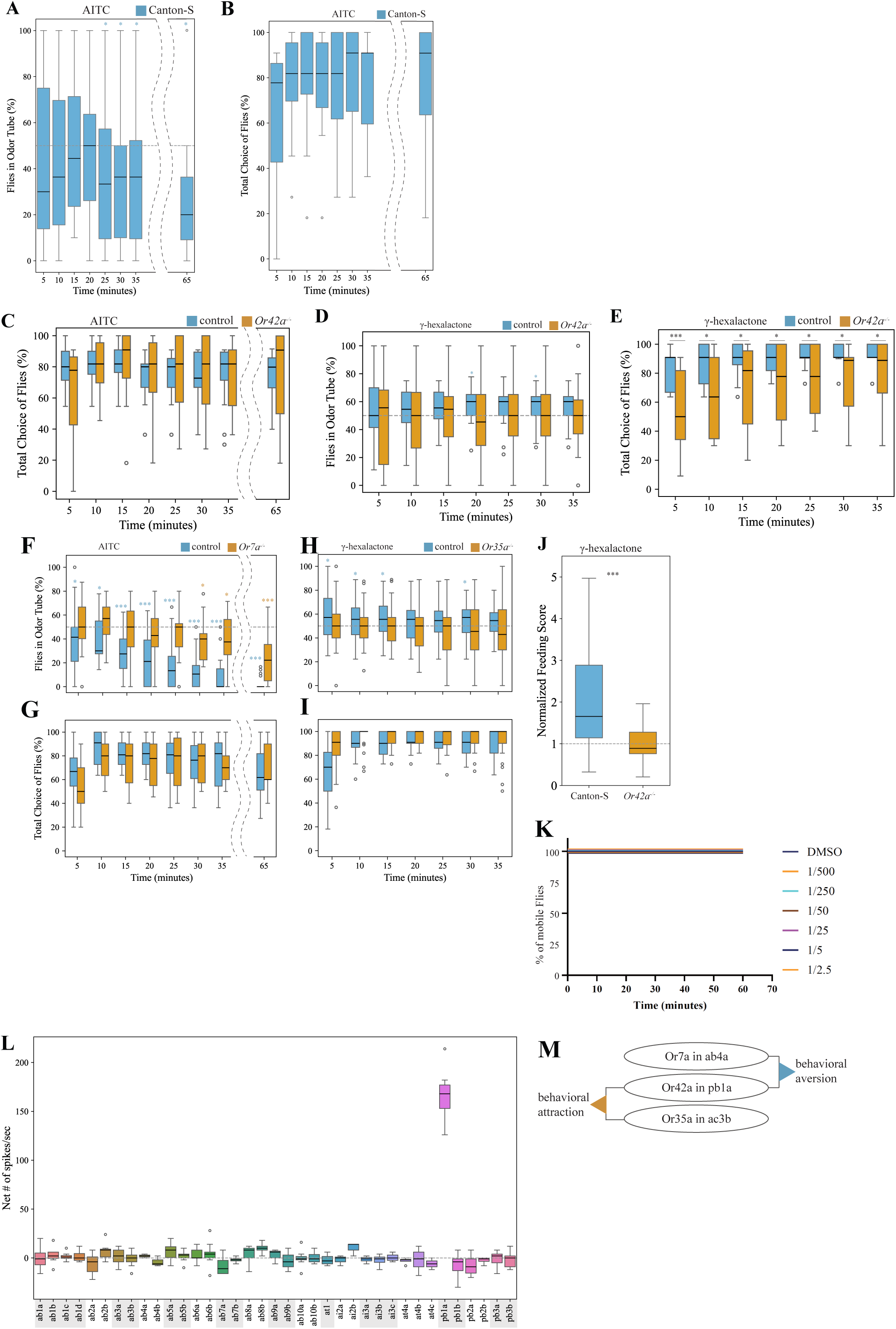
Behavioral data related to Figure 4. Positional olfactory assays. **(A-B)** Tests with allyl isothiocyanate (AITC) 1:500 vol/vol vs mineral oil (as in Figure 4A) using Canton-S flies (n=19); flies preferred the odorless tube from minute 25 onwards (one sample signed rank tests against median=50%, *p<0.05 in all cases; **A**). Total choice of Canton-S flies calculated as: [(# of flies in the odorless tube + # of flies in the odorous tube) / (# of flies released) *100)]; **B**. **(C)** Total choice of genetic background control (line # 68384) and *Or42a-/-* mutants (n=15 for each genotype). Similar percentages of control flies and *Or42a-/-* mutants choose one or the other tube at all time points, indicating that all fly genotypes are similarly active in presence of AITC volatiles (Mann-Whitney U tests, p>0.05 for all time points). **(D-E)** Tests using γ-hexalactone 1:10 vol/vol vs the solvent control using genetic background control flies and *Or42a-/-* mutants. Control flies choose the odorous tube at 20 and 30 minutes (n=19, *p<0.05, one-sample signed rank tests), while mutants distributed randomly at all time points (n=19, p>0.05 in all cases); **D**. Additionally, the total choice of control flies was larger than that of mutant flies at all time points, suggesting that the presence of the odor increases exploration activity in control flies (Mann-Whitney U tests, *p<0.05, ***p<0.005); **E**. Canton-S flies also showed attraction to this odor (at 5, 10, and 15 minutes, n=12; one-sample signed rank tests, p<0.05 at all those three times, data shown in Supplementary File 4). **(F-G)** Tests with *Or7a-/-* (n=15) and genetic background control flies (n=18) using AITC 1:500 vol/vol vs mineral oil. Control flies avoided the AITC tube at all time points, while mutants only avoided at 30 minutes and onwards (one-sample signed rank tests, 1-tail, *p<0.05, ****p<0.001); **F**. The total choice of flies was not different between genotypes (p>0.05 at all time points); **G**. **(H-I)** Tests with *Or35a-/-* (n=29) and genetic control flies (n=27) using γ-hexalactone 1:10 vol/vol vs the solvent control. Control flies choose the odorous tube at various time points (*p<0.05, ***p<0.001, one sample signed rank tests, 1 tail), while mutants always distributed at random (p>0.05 in all cases); **H**. **I:** Total choice of control and Or35a-/- mutants; differences between genotypes were not statistically different (Mann-Whitney U tests, p>0.05 at all time points). In all cases (panels **C-I,** Figure 4A) mutants of the various genotypes used were assayed in parallel with the genetic background control line and therefore, control data was obtained from different subsets of flies. **(J)** Feeding assay, as in Figure 1E, but using γ-hexalactone (1:50 vol/vol). The dotted line at 50% indicates neither feeding enhancement or repellence. Canton-S flies consumed more in presence than in absence of γ-hexalactone (one-sample signed rank test on normalized data against expected median=1, p<0.001, n=25), and *Or42a-/-* mutants instead consumed less in presence of the odor; p<0.001, n=24). (**K)** Mobility assay, as in Figure 2, but using γ-hexalactone at various concentrations from 1:500 to 1:2.5 vol/vol. n=10 for each condition (solvent and concentration). Flies remained active in the presence of this volatile. **(L)** Single sensillum recordings from all *D. melanogaster* antennae (ab) and palp basiconic (pb) sensilla upon stimulation with γ-hexalactone 1:100 vol/vol. Only pb1a was activated by γ-hexalactone (n= 3-6 OSNs). **(M)** Activation of Or42a, in combination with distinct olfactory channels, is involved in mediating both olfactory repellence to AITC (light blue arrow) and attraction to γ-hexalactone (orange arrow). Activation of Or42a in combination with Or7a or O35a likely mediates repellence and attraction, respectively.

**Figure Supplement 3:**
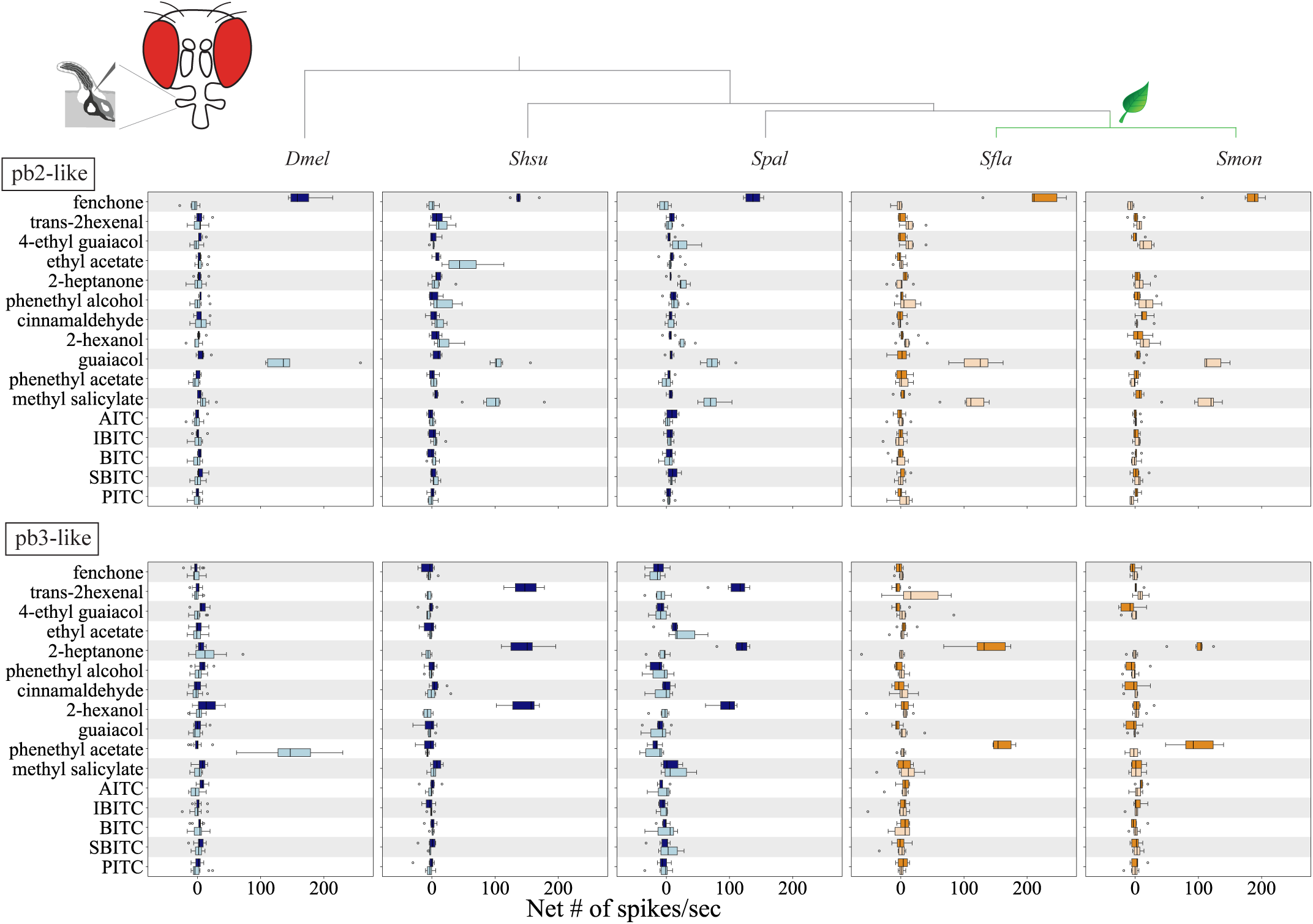
Functional characterization of *D. melanogaster* and *Scaptomyza* species maxillary palp OSNs housed in the three palp basiconic sensilla types. Single sensillum recordings from maxillary palp OSNs (pb2, pb2-like, pb3, and pb3-like) of *D. melanogaster, S. hsui, S. pallida, S. flava,* and *S. montana*. Stimuli (1:100 vol/vol) included diagnostic chemicals used to identify Ors in *D. melanogaster* (see Methods), fruit volatiles, green leaf volatiles, and Brassicales plant-derived isothiocyanates (ITCs) (n=6-9 from 3-4 animals/species). Pb and pb-like sensilla housed two OSNs, labeled “a” (darker color) and “b” (lighter color). Their response profiles and stereotyped locations within the palp support the classification of *Scaptomyza* sensilla into three types, pb1-like, pb2-like, and pb3-like (see also Figure 5). Methyl salicylate (a *S. flava* volatile attractant in natural settings, Orre et al. 2010) activated all *Scaptomyza* pb2b-like sensilla but not *D. melanogaster* pb2b (first row). The response profiles of *Scaptomyza* pb3-like OSNs were different from those of *D. melanogaster* pb3, likely because the homologs of *Or59c* and *Or85d*, which are respectively expressed in *D. melanogaster* pb3a and p3b, are unidentified in the genomes of *Scaptomyza* (Goldman-Huertas et al. 2015). The odorant response profiles of OSNs in pb3-like sensilla were more similar between the more distantly related species *S. hsui* and *S. pallida,* than between the more closely related *S. pallida* and *S. flava,* or *S. pallida* and *S. montana.* For example, *trans*-2-hexenal activated these OSNs in *S. hsui* and *S. pallida* (but not in *S. flava* or *S. montana*), while phenethyl acetate activated *S. flava* and *S. montana* OSNs (but not *S. hsui* or *S. pallida*). AITC: allyl ITC, IBITC: isobutyl ITC, BITC: butyl ITC, SBITC: *sec*-butyl ITC, PITC: phenethyl ITC.

**Figure Supplement 4:**
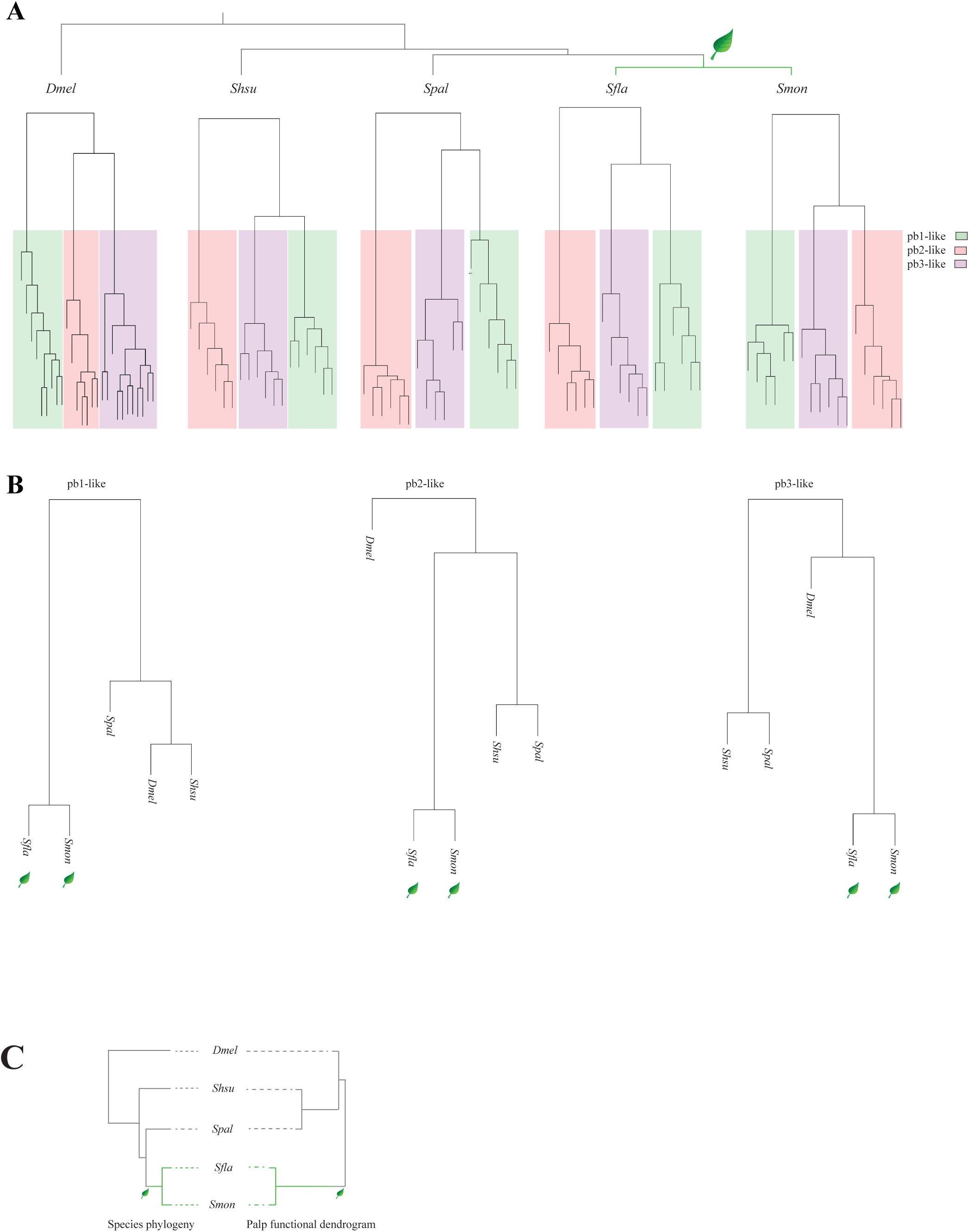
Hierarchical cluster analysis of maxillary palp sensilla based on odorant response profiles across species. **(A)** Hierarchical clusters were constructed using odorant response data from single sensillum recordings of basiconic palp sensilla from *D. melanogaster, S. hsui, S. pallida, S. flava,* and *S. montana* using R studio v1.4.1717. A terminal node corresponds to a single recording from the individual sensilla; n=6-8 sensilla from 3-4 animals. **(B)** Hierarchical clusters were created for each sensilla type, including pb1-like, pb2-like, and pb3-like, using the odorant response (control-subtracted net number of spikes) average from each species as inputs. **(C)** The species phylogeny (left: Kim et al., 2021) and palp functional dendrogram (right) highlight the relevance of niche differences in shaping odorant responses. The palp functional dendrogram was generated using odorant response averages from each species.

**Figure Supplement 5.**
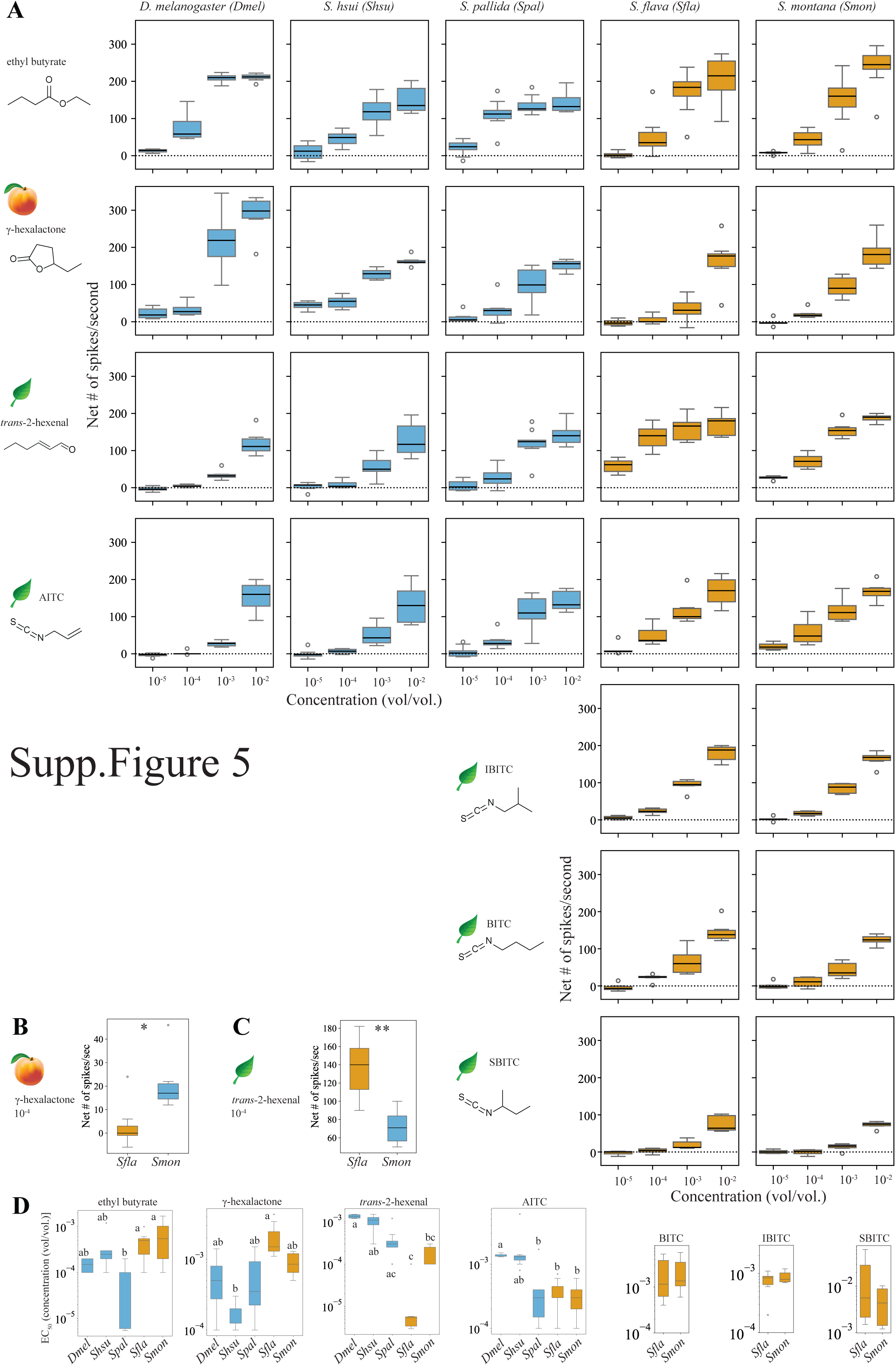
Responses of OSNs from *D. melanogaster* pb1a and *Scaptomyza* pb1a-like sensilla to stimulation with various odorant concentrations. (A) Responses of pb1a OSNs across species upon stimulation with various concentrations (ranging from 1:10^-2^ to 10^-5^ vol/vol) of ethyl butyrate, γ-hexalactone, *trans*-2-hexenal, allyl isothiocyanate (AITC), isobutyl isothiocyanate (IBITC), butyl isothiocyanate (BITC) and *sec*-butyl isothiocyanate (SBITC; n=6-8 sensilla from 3-4 animals/species). The chemical structures of each compound are shown on the left. IBITC, BITC, and SBITC were only tested in *S. flava* and *S. montana* pb1a, as BITC at 10^-2^ vol/vol failed to activate OSNs in pb1a from the two microbe-feeding species (Figure 5). **(B-C)** Responses of *S. flava* and *S. montana* pb1a OSNs upon stimulation with γ-hexalactone 10^-4^ vol/vol **(B)** and the green leaf volatile *trans*-2-hexenal 10^-4^ vol/vol **(C)**; n=6-7 sensilla from 3 animals/species. Mann-Whitney U tests: *p<0.05, **p<0.01. **(D)** Half maximal effective concentrations (EC_50_) for each fly species and odorant (see Methods for calculations of EC_50_). Kruskal-Wallis tests followed by Dunn’s multiple comparisons. Different letters indicate significant differences between species (p-values for ethyl butyrate: *S. pallida* vs *S. flava* p<0.05, *S. pallida* vs *S. montana* p<0.01; for γ-hexalactone: p<0.001; for *trans*-2-hexenal: *D. melanogaster* vs *S. montana* p<0.05, *S. hsui* vs *S. flava* p <0.01, *D. melanogaster* vs *S. flava* p<0.0001; for AITC: p<0.05). The EC_50s_ of *S. flava* and *S. montana* to BITC, IBITC and SBITC were not statistically different (Mann-Whitney U tests, p>0.05). Overall, pb1a OSNs of microbe-feeding species exhibited greater sensitivity to γ-hexalactone, whereas Brassicales specialists showed higher sensitivity to *trans*-2-hexenal and AITC. *S. flava* and *S. montana* showed similar sensitivities to all four ITCs tested.

**Figure Supplement 6:**
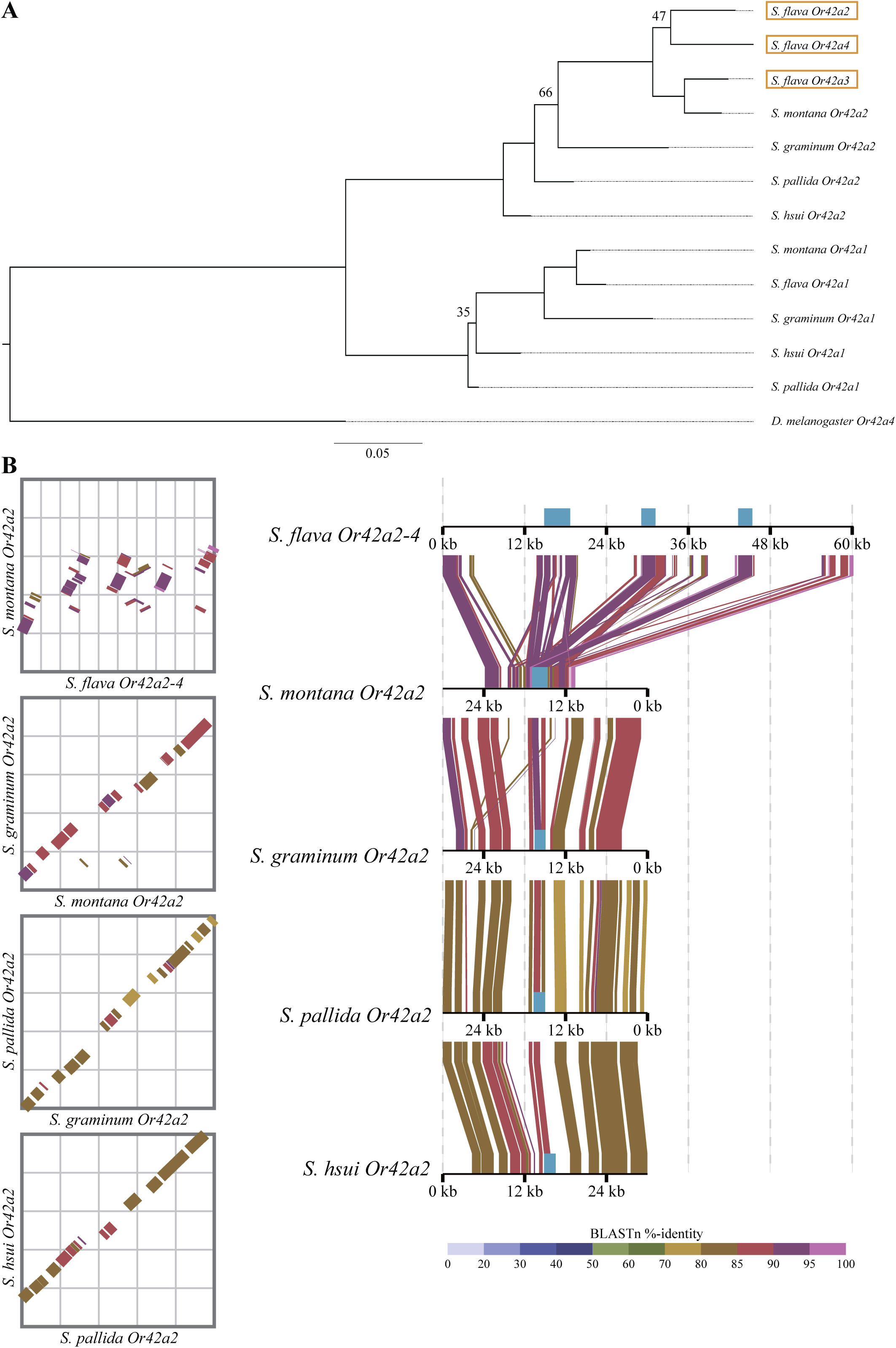
Or42a gene tree constructed by RAxML, and syntenic maps and dot plots visualized by DiGAlign. **(A)** Maximum-likelihood gene tree with 1000 bootstrap cycles using *S. hsui Or42a1*, *S. pallida Or42a1*, *S. graminum Or42a1*, *S. montana Or42a1*, *S. flava Or42a1*, *S. hsui Or42a2*, *S. pallida Or42a2*, *S. graminum Or42a2*, *S. flava Or42a2*, *S. montana Or42a*, *S. flava Or42a2*, *S. flava Or42a3*, *S. flava Or42a4*, and *D. melanogaster Or42a* as an outgroup, generated by RAxML (v8.2.19) (Stamatakis 2014) after alignment by MAFFT v7 (Katoh and Standley 2013). Bootstrap values are shown on the branches when the values were lower than 80. The tree was displayed by FigTree v1.4.3 (Rambaut 2009). **(B)** Dot plots (left) and synteny maps (right) were generated for the *Or42a2-Or42a4* genomic region across four species pairs: *S. flava*-*S. montana*, *S. montana*-*S. graminum*, *S. graminum*-*S. pallida*, and *S. pallida*-*S. hsui*. A 60,199-nt region was analyzed in *S. flava*, and 30,000-nt regions were analyzed in the other four species. Genomes of *S. montana, S. graminum,* and *S. pallida* were reversed-aligned for visualization. Each genomic region is color-coded according to BLASTn percent identity (top left). Cyan rectangles mark the span from the start to the end of *Or42a* homologs. *S. flava* has three copies of *Or42a2* species-specific, separated by intronic intervals that are mostly not conserved in the other species. In contrast, the remaining *Scaptomyza* species showed conserved synteny at the *Or42a2* locus.

**Figure Supplement 7:**
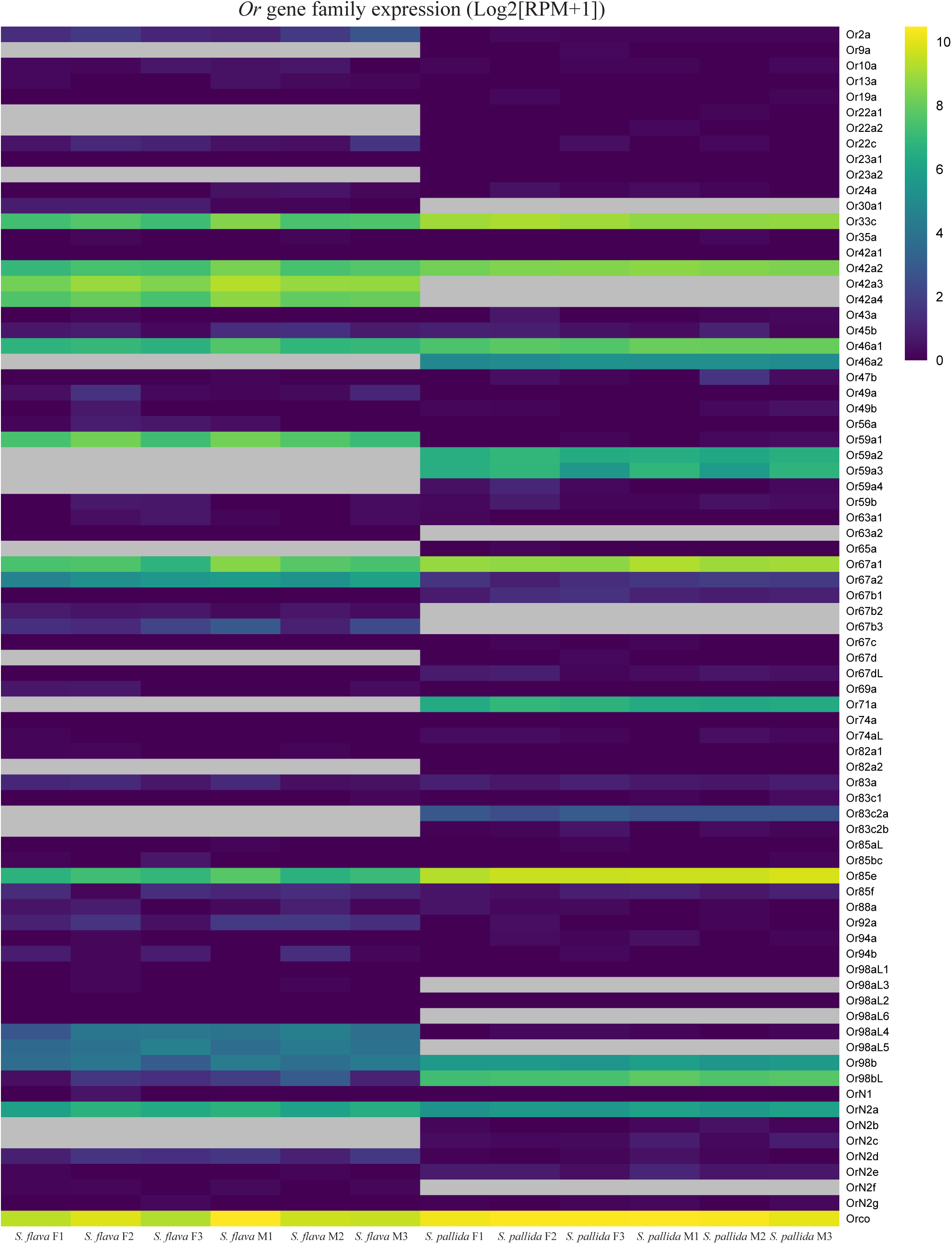
Maxillary palp RNA seq of *S. pallida* and *S. flava Or*s. The heatmap displays Log2[RPM+1] values of *Or*s expression in the maxillary palps of female (F) and male (M) *S. flava* and *S. pallida* (n=3/sex). Each column represents a replicate (e.g. *S. flava* F1: replicate #1 of female *S. flava*, *S. flava* M1: replicate #1 of male *S. flava*). Gray boxes indicate that the corresponding genes were unidentified in the genomes of the species. Only one copy of Or42a (Or42a2) is expressed in *S. pallida*, while *S. flava* expresses three copies. We confirmed the expression of homologs *Or71a* (expressed in OSNs of *D. melanogaster* pb1b), and of *Or33c/Or85e* and *Or46a* (expressed in OSNs of *D. melanogaster* pb2a and pb2b) in both *S. pallida* and *S. flava* maxillary palps. *Or85d* and *Or59c* were respectively expressed in *D. melanogaster* pb3a and pb3b OSNs but the homologs were unidentified in the genomes of *Scaptomyza* (Goldman-Huertas et al., 2015). Instead, *Or59a1, Or67a1, Or67a2,* and *OrN2a* were strongly expressed in OSNs of both *S. pallida* and *S. flava,* suggesting that these homologs may be expressed in pb3a or pb3b.

**Figure Supplement 8:**
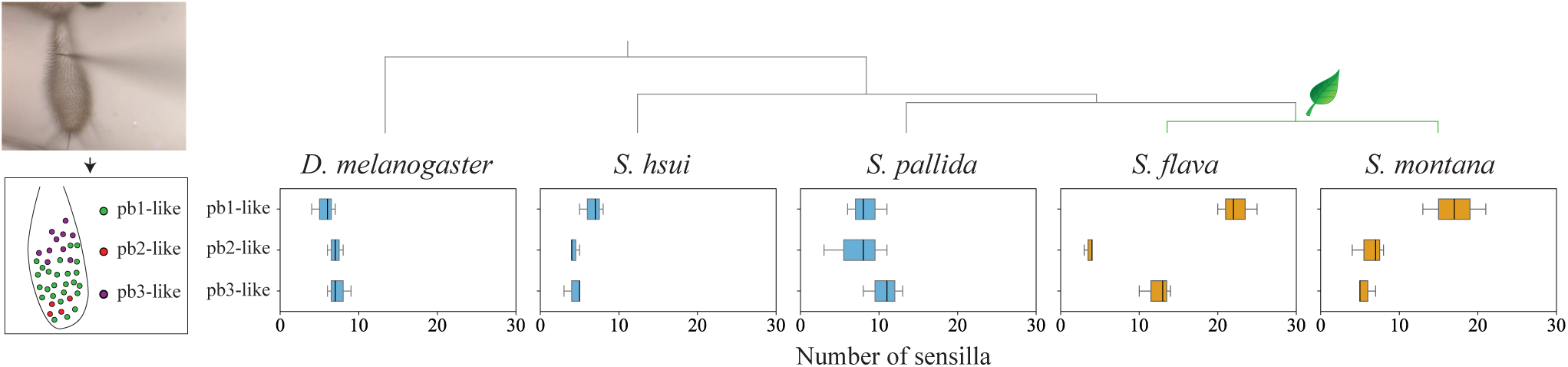
Over-representation of pb1-like sensilla in mustard plant specialists. Picture of single sensillum recording from *S. flava* maxillary palp sensilla (top left) and a representative anatomical mapping of *S. flava* palp sensilla (bottom left) obtained using diagnostic chemicals. Green: pb1-like, red: pb2-like, and magenta: pb3-like. Boxplots in the graph (right) represent the number of each sensilla type in *D. melanogaster* and the four *Scaptomyza* species (n=3 animals/species; see Figure Supplement 9 for individual maps). Maxillary palp pb1 sensilla are over-represented in *S. flava* and *S. montana*, the two mustard specialists.

**Figure Supplement 9:**
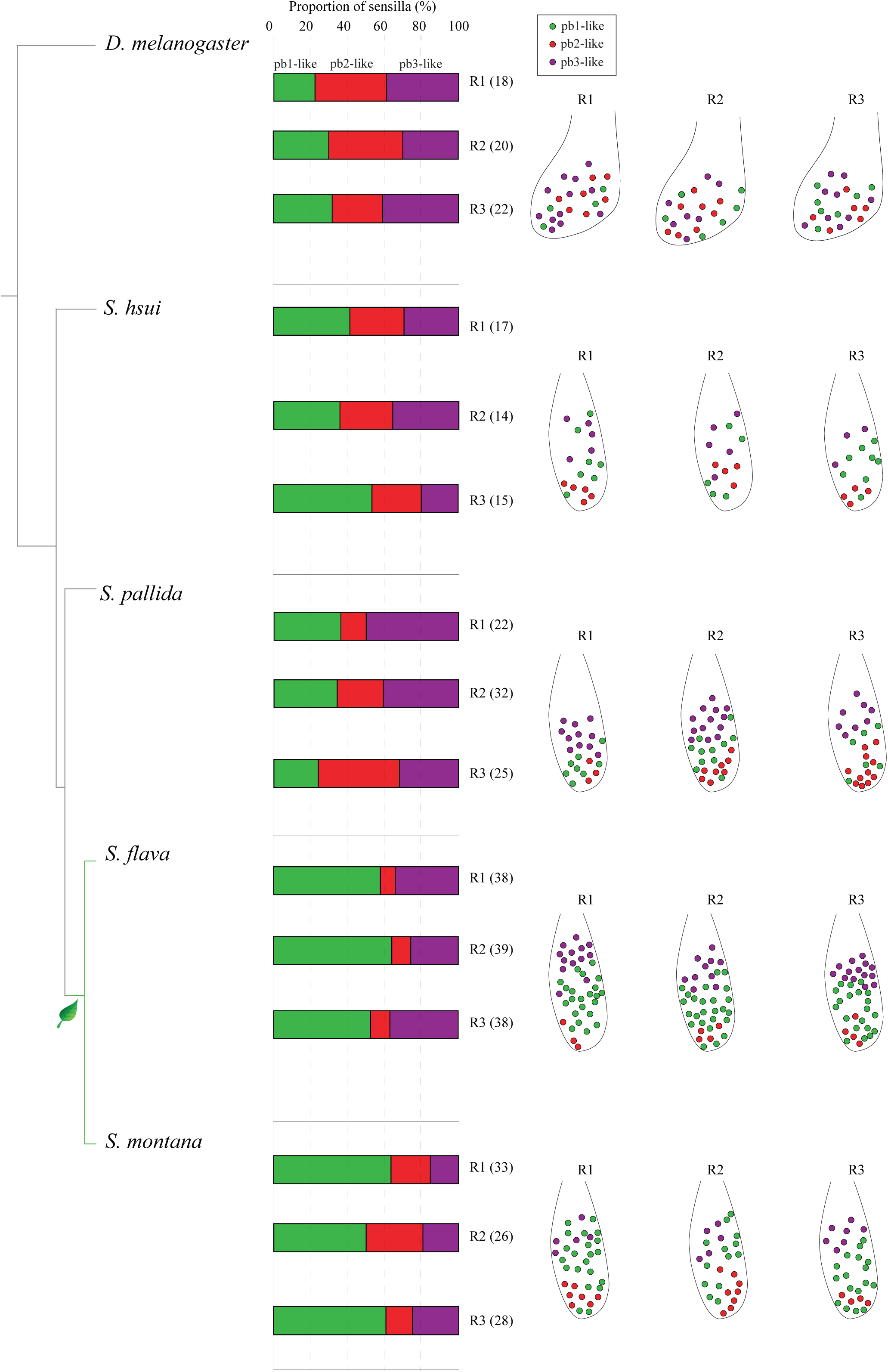
Location and percentages of the three different maxillary palp basiconic (pb) sensilla across species. Classification of maxillary palp pb sensilla using single sensillum recordings and stimulation with diagnostic odorants. Data represents the proportion of pb1 and pb1-like (green), pb2 and pb2-like (red), and pb3 and pb3-like (magenta) in *D. melanogaster, S. hsui, S. pallida, S. flava,* and *S. montana*. The schematic positions of pb sensilla in each of the three replicates (R1, R2, and R3) from each species are shown on the right. The total number of sensilla identified in each animal is indicated between parentheses.

**Figure Supplement 10:**
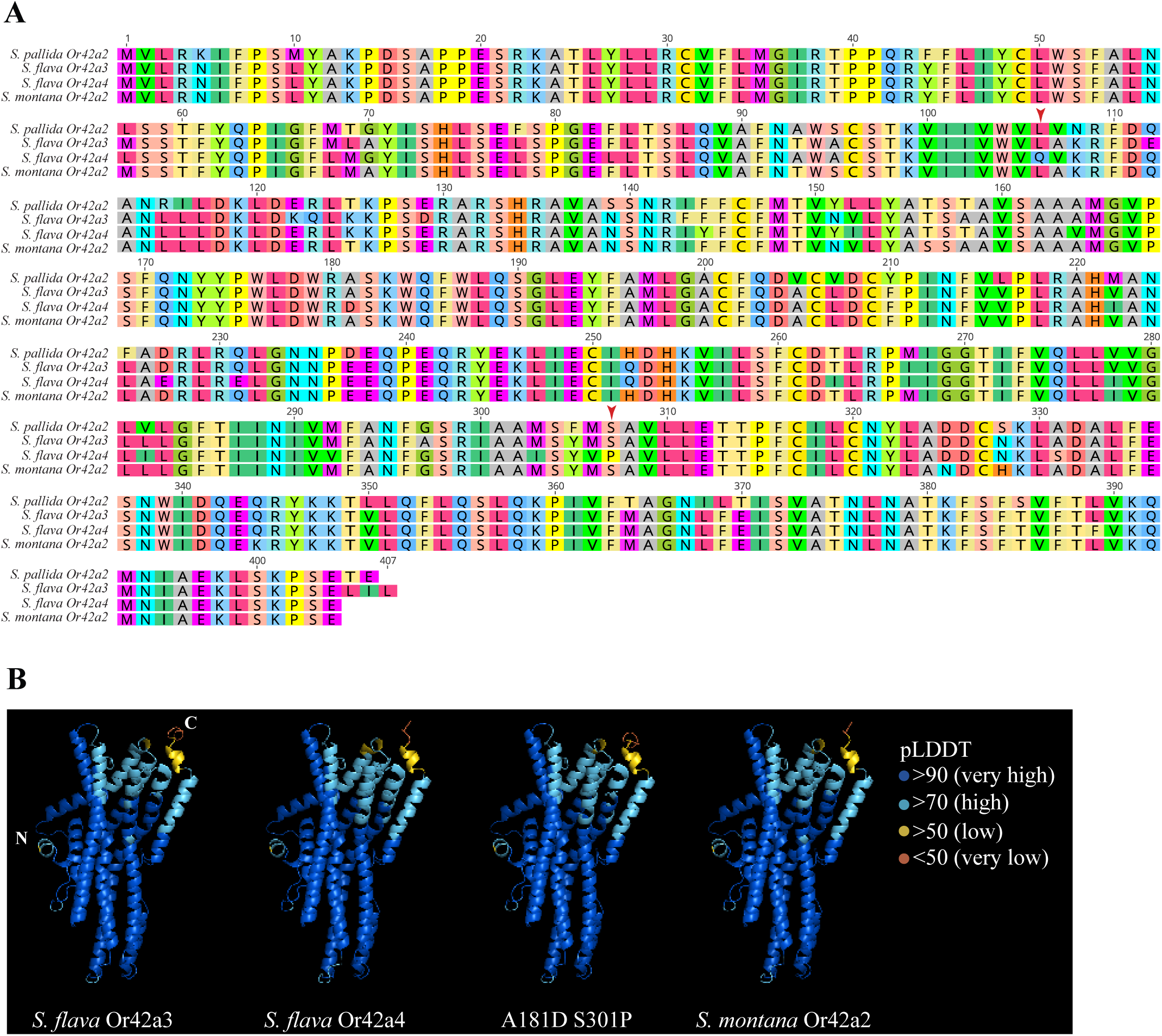
Alignment of 1D and 3D structures of amino acids of Or42a. **(A)** The 1D alignment of amino acids of *S. pallida* Or42a2, *S. flava* Or42a3, *S. flava* Or42a4, and *S. montana* Or42a2. Red arrowheads on top denote the sites where site-directed-mutagenesis was performed to generate the A181D S301P chimera. **(B)** The 3D structures of *S. flava* Or42a3, *S. flava* Or42a4, A181D S301P, and *S. montana* Or42a2 were predicted using AlphaFold2. The colors of the amino acids indicate the predicted local distance difference test (pLDDT) scores: blue (>90), cyan (>70), yellow (> 50), and orange (<50). We confirmed that the scores for these non-overlapping sites were above 70, indicating that the 3D predictions of these sites are reliable.

**Figure Supplement 11:**
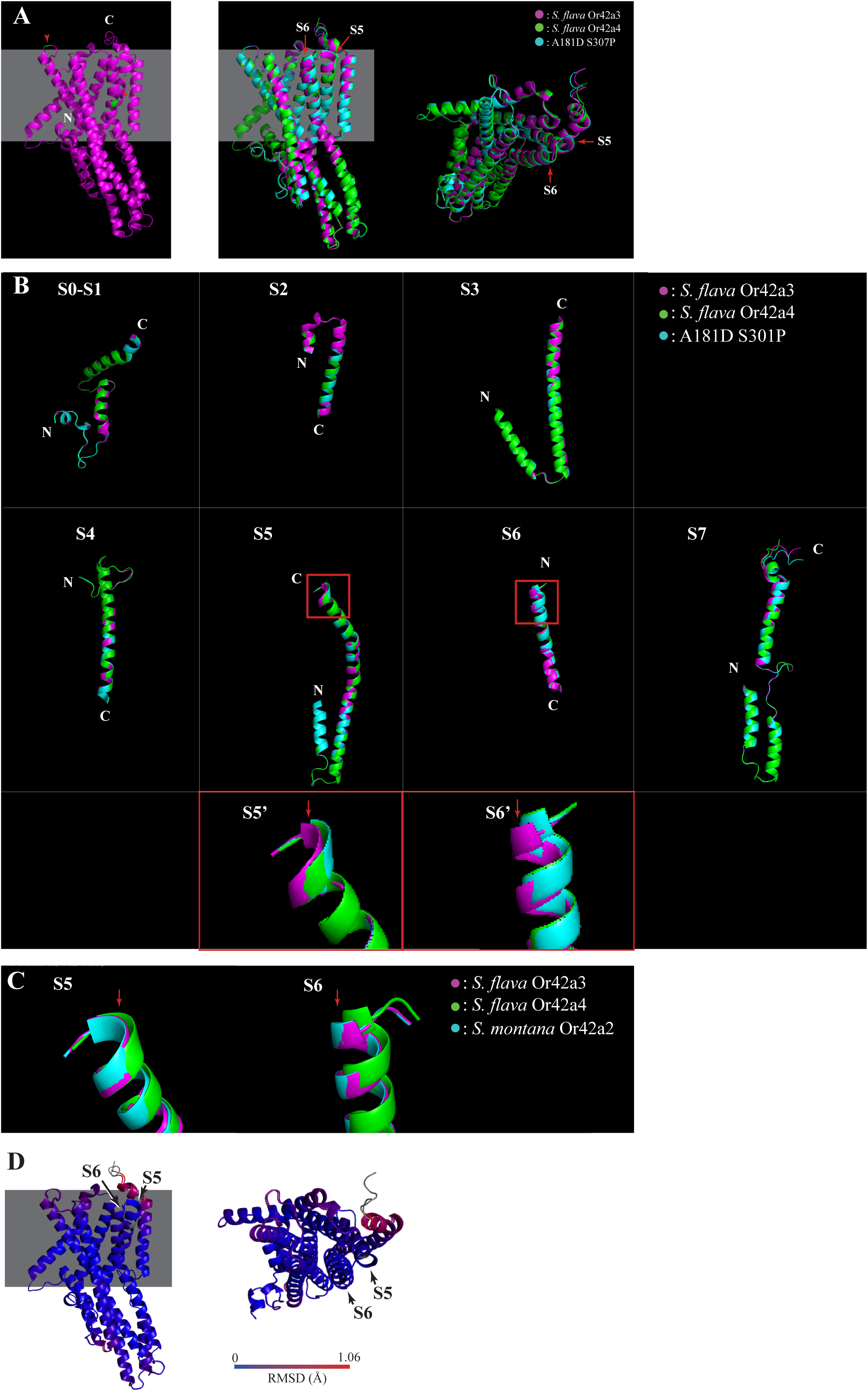
Screening of candidate amino acids by AlphaFold2 3D prediction. **(A)** Lateral view of the 3D structure of A181D S301P predicted by AlphaFold2 (left). *S. flava* Or42a3-derived amino acids are indicated in magenta, and the substitutions with *S. flava* Or42a4-derived amino acids are represented in green. The middle and right panels respectively display side and top views of alignment of the predicted structures of *S. flava* Or42a3, *S. flava* Or42a4, and A181D S301P. The extracellular region at S5 and S6 (red arrows) highlights the local structural differences between *S. flava* Or42a3 and Or42a4 where the A181D, S301P overlaps with the local structure of *S. flava* Or42a4. **(B)** The 3D predictions of *S. flava* Or42a3 (magenta), *S. flava* Or42a4 (green), and A181D S301P (cyan) were aligned using PyMol 2.5.3. Red rectangles highlight the sites of local structural variance between *S. flava* Or42a3 and Or42a4. The A181D S301P shows a closer alignment with Or42a4. S5’ and S6’ are enlarged views of S5 and S6, respectively. **(C)** The enlarged views of S5 and S6 in the alignment of 3D predictions of *S. flava* Or42a3 (magenta), *S. flava* Or42a4 (green), and *S. montana* Or42a2 (cyan). **(D)** The 3D alignment of *S. flava* Or42a3 and *S. montana* Or42a2 predicted by AlphaFold2. The root mean square deviation (RMSD) is visualized with a color gradient.

**Figure Supplement 12:**
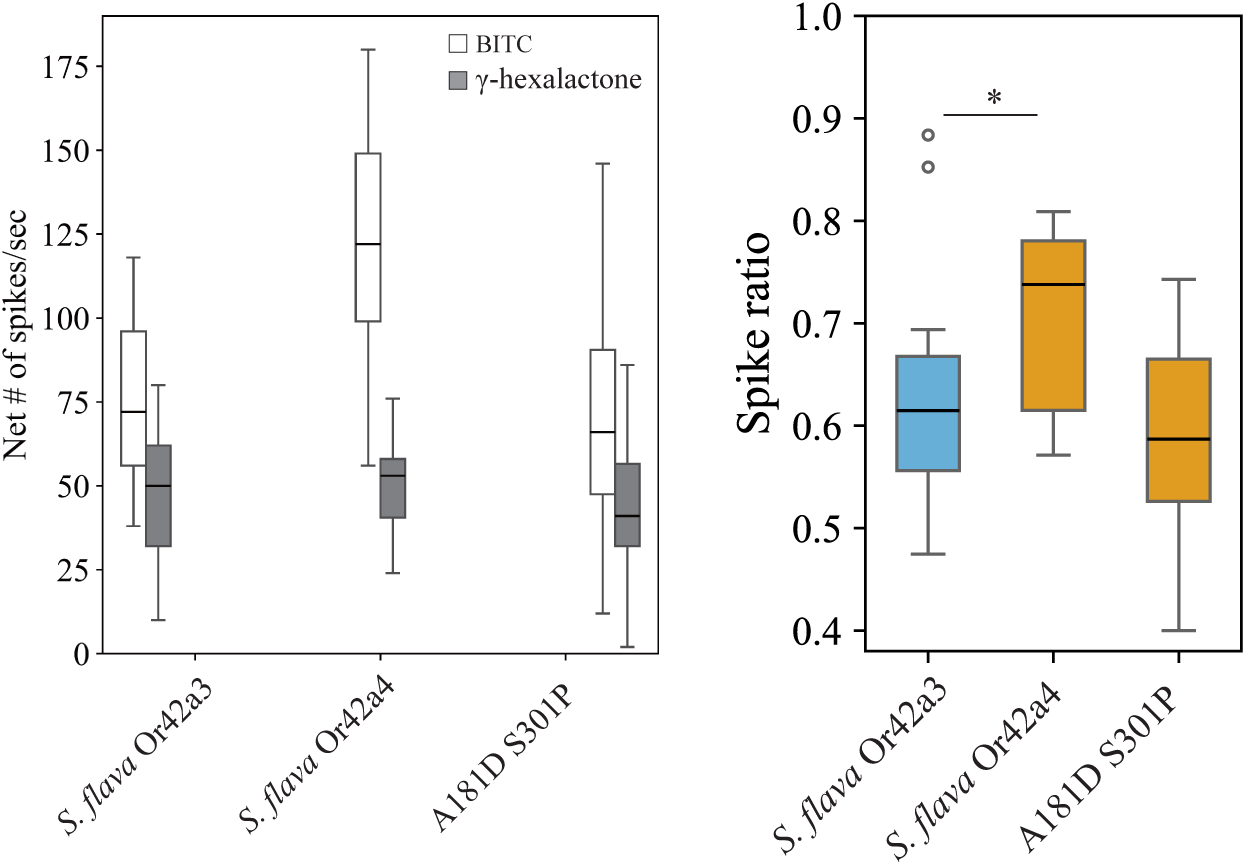
Responses of OSNs expressing homozygous Or42a to stimulation with butyl isothiocyanate and γ-hexalactone. Left: responses of OSNs expressing homozygous *S. flava* Or42a3, *S. flava* Or42a4, or A181D S307P (*UAS-Or42a; Or67d ^Gal4^*) to stimulation with butyl isothiocyanate (BITC, white) and γ-hexalactone (grey) (n=16-30 from 5-10 animals). Right: ratio between responses to BITC and the sum of the responses evoked by stimulation with BITC and γ-hexalactone. Kruskal-Wallis followed by Dunn’s multiple comparisons, *p<0.05.

**Figure Supplement 13:**
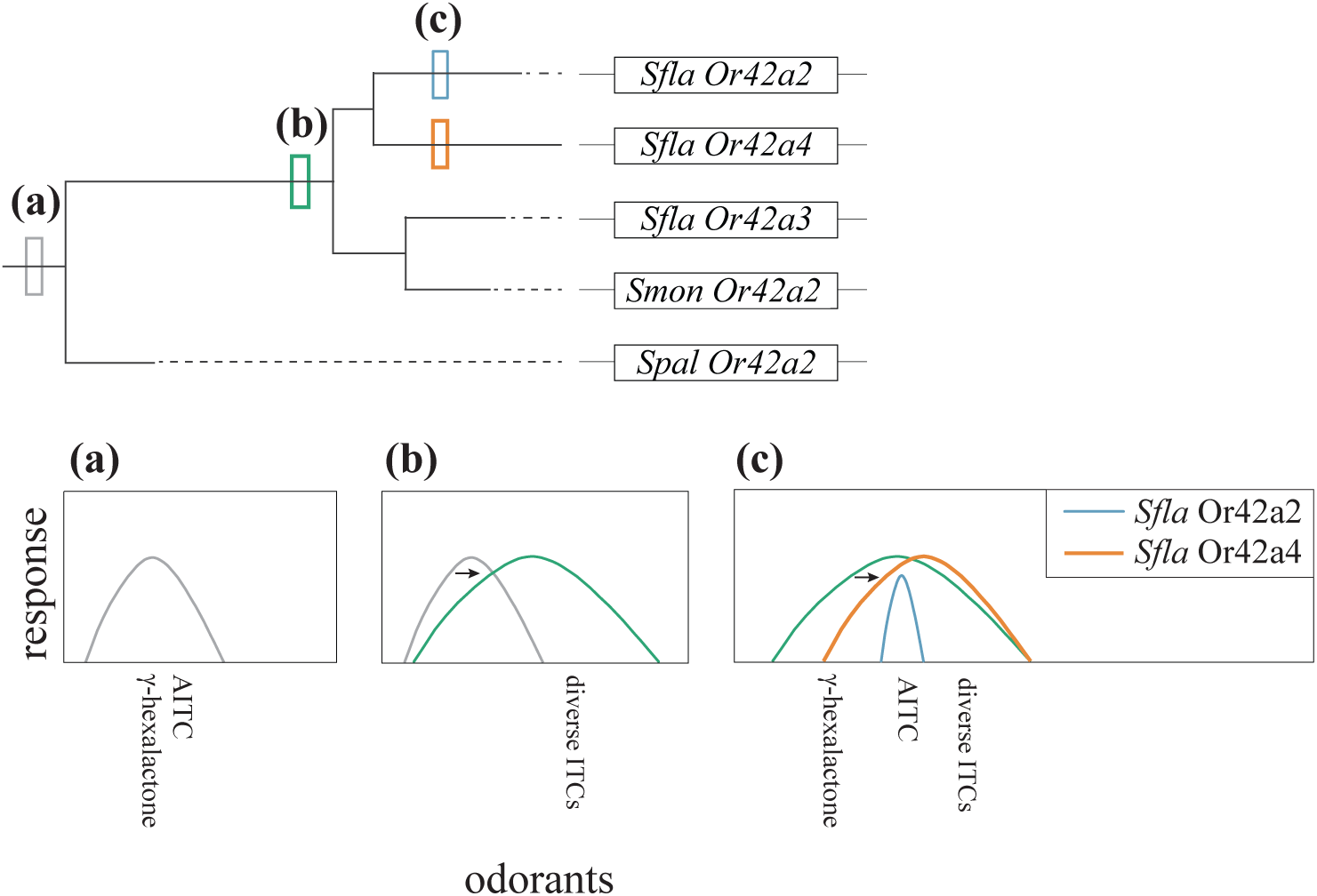
A model for the evolution of Or42a in *S. flava, S. montana* and *S. pallida*. The evolution of Or42a begins with a shift in the ligand specificity of an ancestral Or42a (a), which was tuned to fruit-borne odors such as γ-hexalactone and limited ITCs (i.e. AITC), and later it broadened to detect a wider range of ITCs (b). Subsequent gene triplication of *S. flava Or42a* resulted in three paralogous genes (c: *S. flava Or42a2, S. flava Or42a3,* and *S. flava Or42a4*). The lineage leading to *S. flava* Or42a2 and *S. flava* Or42a4 experienced a reduction in sensitivity to γ-hexalactone (arrow pointing to the right in the bottom rightmost panel), while both *S. flava* Or42a3 and *S. montana* Or42a2 retained sensitivity to γ-hexalactone.

## REFERENCES

1. Aguilar JM, Gloss AD, Suzuki HC, Verster KI, Singhal M, Hoff J, Grebenok R, Nabity PD, Behmer ST, Whiteman NK. 2024. Insights into the evolution of herbivory from a leaf-mining fly. Ecosphere [Internet] 15. Available from: https://esajournals.onlinelibrary.wiley.com/doi/10.1002/ecs2.4764

2. Ahuja I, Rohloff J, Bones AM. 2010. Defence mechanisms of Brassicaceae: implications for plant-insect interactions and potential for integrated pest management. A review. Agron. Sustain. Dev. 30:311–348.

3. Al-Anzi B, Tracey WD Jr, Benzer S. 2006. Response of Drosophila to wasabi is mediated by painless, the fly homolog of mammalian TRPA1/ANKTM1. Curr. Biol. 16:1034–1040.

4. Anders S, Pyl PT, Huber W. 2015. HTSeq--a Python framework to work with high-throughput sequencing data. Bioinformatics 31:166–169.

5. Auer TO, Khallaf MA, Silbering AF, Zappia G, Ellis K, Álvarez-Ocaña R, Arguello JR, Hansson BS, Jefferis GSXE, Caron SJC, et al. 2020. Olfactory receptor and circuit evolution promote host specialization. Nature 579:402–408.

6. Becher PG, Bengtsson M, Hansson BS, Witzgall P. 2010. Flying the fly: long-range flight behavior of Drosophila melanogaster to attractive odors. J. Chem. Ecol. 36:599–607.

7. Bell L, Oloyede OO, Lignou S, Wagstaff C, Methven L. 2018. Taste and flavor perceptions of glucosinolates, isothiocyanates, and related compounds. Mol. Nutr. Food Res. 62:e1700990.

8. Benton R, Himmel NJ. 2023. Structural screens identify candidate human homologs of insect chemoreceptors and cryptic Drosophila gustatory receptor-like proteins. Elife 12:e85537.

9. Bernays E, Chapman R. 1987. The evolution of deterrent responses in plant-feeding insects. In: Proceedings in Life Sciences. New York, NY: Springer New York. p. 159–173.

10. Bischof J, Maeda RK, Hediger M, Karch F, Basler K. 2007. An optimized transgenesis system for Drosophila using germ-line-specific phiC31 integrases. Proc. Natl. Acad. Sci. U. S. A. 104:3312–3317.

11. de Bruyne M, Clyne PJ, Carlson JR. 1999. Odor coding in a model olfactory organ: the Drosophila maxillary palp. J. Neurosci. 19:4520–4532.

12. Butterwick JA, Del Mármol J, Kim KH, Kahlson MA, Rogow JA, Walz T, Ruta V. 2018. Cryo-EM structure of the insect olfactory receptor Orco. Nature 560:447–452.

13. Chen S, Zhou Y, Chen Y, Gu J. 2018. fastp: an ultra-fast all-in-one FASTQ preprocessor. Bioinformatics 34:i884–i890.

14. Chen Y-CD, Dahanukar A. 2020. Recent advances in the genetic basis of taste detection in Drosophila. Cell. Mol. Life Sci. 77:1087–1101.

15. Chou PY, Fasman GD. 1978. Prediction of the secondary structure of proteins from their amino acid sequence. Adv. Enzymol. Relat. Areas Mol. Biol. 47:45–148.

16. Couto A, Alenius M, Dickson BJ. 2005. Molecular, anatomical, and functional organization of the Drosophila olfactory system. Curr. Biol. 15:1535–1547.

17. Crowley-Gall A, Date P, Han C, Rhodes N, Andolfatto P, Layne JE, Rollmann SM. 2016. Population differences in olfaction accompany host shift in Drosophila mojavensis. Proc. Biol. Sci. [Internet] 283. Available from: https://www.ncbi.nlm.nih.gov/pmc/articles/PMC5013806/

18. Cuellar-Nuñez ML, Luzardo-Ocampo I, Lee-Martínez S, Larrauri-Rodríguez M, Zaldívar-Lelo de Larrea G, Pérez-Serrano RM, Camacho-Calderón N. 2022. Isothiocyanate-Rich Extracts from Cauliflower (Brassica oleracea Var. Botrytis) and Radish (Raphanus sativus) Inhibited Metabolic Activity and Induced ROS in Selected Human HCT116 and HT-29 Colorectal Cancer Cells. Int. J. Environ. Res. Public Health [Internet] 19. Available from: 10.3390/ijerph192214919

19. Dekker T, Ibba I, Siju KP, Stensmyr MC, Hansson BS. 2006. Olfactory shifts parallel superspecialism for toxic fruit in Drosophila melanogaster sibling, D. sechellia. Curr. Biol. 16:101–109.

20. Del Mármol J, Yedlin MA, Ruta V. 2021. The structural basis of odorant recognition in insect olfactory receptors. Nature 597:126–131.

21. Dobin A, Davis CA, Schlesinger F, Drenkow J, Zaleski C, Jha S, Batut P, Chaisson M, Gingeras TR. 2013. STAR: ultrafast universal RNA-seq aligner. Bioinformatics 29:15–21.

22. Dobler S, Dalla S, Wagschal V, Agrawal AA. 2012. Community-wide convergent evolution in insect adaptation to toxic cardenolides by substitutions in the Na,K-ATPase. Proc. Natl. Acad. Sci. U. S. A. 109:13040–13045.

23. Dobritsa AA, van der Goes van Naters W, Warr CG, Steinbrecht RA, Carlson JR. 2003. Integrating the molecular and cellular basis of odor coding in the Drosophila antenna. Neuron 37:827–841.

24. Dweck HK, Ebrahim SA, Khallaf MA, Koenig C, Farhan A, Stieber R, Weißflog J, Svatoš A, Grosse-Wilde E, Knaden M, et al. 2016. Olfactory channels associated with the Drosophila maxillary palp mediate short- and long-range attraction. Elife [Internet] 5. Available from: https://www.ncbi.nlm.nih.gov/pmc/articles/PMC4927298/

25. Dweck HKM, Carlson JR. 2020. Molecular logic and evolution of bitter taste in Drosophila. Curr. Biol. 30:17–30.e3.

26. Ferreiro MJ, Pérez C, Marchesano M, Ruiz S, Caputi A, Aguilera P, Barrio R, Cantera R. 2017. Drosophila melanogaster white mutant w1118 undergo retinal degeneration. Front. Neurosci. 11:732.

27. Gloss AD, Brachi B, Feldmann MJ, Groen SC, Bartoli C, Gouzy J, LaPlante ER, Meyer CG, Pyon HS, Rogan SC, et al. 2017. Genetic variants affecting plant size and chemical defenses jointly shape herbivory in*Arabidopsis*. bioRxiv [Internet]:156299. Available from: https://www.biorxiv.org/content/10.1101/156299v1

28. Goldman-Huertas B, Mitchell RF, Lapoint RT, Faucher CP, Hildebrand JG, Whiteman NK. 2015. Evolution of herbivory in Drosophilidae linked to loss of behaviors, antennal responses, odorant receptors, and ancestral diet. Proceedings of the National Academy of Sciences 112:3026–3031.

29. Gonzalez F, Witzgall P, Walker WB. 2016. Protocol for heterologous expression of insect odourant receptors in Drosophila. Front. Ecol. Evol. [Internet] 4. Available from: https://www.frontiersin.org/journals/ecology-and-evolution/articles/10.3389/fevo.2016.00024/full

30. Hashimoto Y, Yoshimura M, Huang R-N. 2019. Wasabi versus red imported fire ants: preliminary test of repellency of microencapsulated allyl isothiocyanate against Solenopsis invicta (Hymenoptera: Formicidae) using bait traps in Taiwan. Appl. Entomol. Zool. (Jpn.) 54:193–196.

31. Haverkamp A, Hansson BS, Knaden M. 2018. Combinatorial codes and labeled lines: How insects use olfactory cues to find and judge food, mates, and oviposition sites in complex environments. Front. Physiol. 9:49.

32. Himmel NJ, Moi D, Benton R. 2023. Remote homolog detection places insect chemoreceptors in a cryptic protein superfamily spanning the tree of life. Curr. Biol. 33:5023–5033.e4.

33. Hopkins RJ, van Dam NM, van Loon JJA. 2009. Role of glucosinolates in insect-plant relationships and multitrophic interactions. Annu. Rev. Entomol. 54:57–83.

34. Ibanez S, Gallet C, Després L. 2012. Plant insecticidal toxins in ecological networks. Toxins (Basel) 4:228–243.

35. Iorio R, Celenza G, Petricca S. 2022. Multi-target effects of ß-caryophyllene and carnosic acid at the Crossroads of mitochondrial dysfunction and neurodegeneration: From oxidative stress to microglia-mediated neuroinflammation. Antioxidants (Basel) 11:1199.

36. Jumper J, Evans R, Pritzel A, Green T, Figurnov M, Ronneberger O, Tunyasuvunakool K, Bates R, Žídek A, Potapenko A, et al. 2021. Highly accurate protein structure prediction with AlphaFold. Nature 596:583–589.

37. Kang K, Pulver SR, Panzano VC, Chang EC, Griffith LC, Theobald DL, Garrity PA. 2010. Analysis of Drosophila TRPA1 reveals an ancient origin for human chemical nociception. Nature 464:597–600.

38. Katoh K, Standley DM. 2013. MAFFT multiple sequence alignment software version 7: improvements in performance and usability. Mol. Biol. Evol. 30:772–780.

39. Kim BY, Wang JR, Miller DE, Barmina O, Delaney E, Thompson A, Comeault AA, Peede D, D’Agostino ERR, Pelaez J, et al. 2021. Highly contiguous assemblies of 101 drosophilid genomes. Elife [Internet] 10. Available from: https://pubmed.ncbi.nlm.nih.gov/34279216/

40. Kim SH, Lee Y, Akitake B, Woodward OM, Guggino WB, Montell C. 2010. *Drosophila* TRPA1 channel mediates chemical avoidance in gustatory receptor neurons. Proceedings of the National Academy of Sciences 107:8440–8445.

41. Kurtovic A, Widmer A, Dickson BJ. 2007. A single class of olfactory neurons mediates behavioural responses to a Drosophila sex pheromone. Nature 446:542–546.

42. Larsson MC, Domingos AI, Jones WD, Chiappe ME, Amrein H, Vosshall LB. 2004. Or83b encodes a broadly expressed odorant receptor essential for Drosophila olfaction. Neuron 43:703–714.

43. Levitt M. 1978. Conformational preferences of amino acids in globular proteins. Biochemistry 17:4277– 4285.

44. Lichtenstein EP, Morgan DG, Mueller CH. 1964. Insecticides in Nature, Naturally Occurring Insecticides in Cruciferous Crops. J. Agric. Food Chem. 12:158–161.

45. Lin C-C, Prokop-Prigge KA, Preti G, Potter CJ. 2015. Food odors trigger Drosophila males to deposit a pheromone that guides aggregation and female oviposition decisions. Elife 4:e08688.

46. Linz J, Baschwitz A, Strutz A, Dweck HKM, Sachse S, Hansson BS, Stensmyr MC. 2013. Host plant-driven sensory specialization in Drosophila erecta. Proc. Biol. Sci. 280:20130626.

47. Liu X-L, Zhang J, Yan Q, Miao C-L, Han W-K, Hou W, Yang K, Hansson BS, Peng Y-C, Guo J-M, et al. 2020. The molecular basis of host selection in a Crucifer-specialized moth. Curr. Biol. 30:4476–4482.e5.

48. MacLeod AJ, MacLeod G, Reader G. 1989. Evidence for the occurrence of butyl- and isobutylglucosinolates in seeds of Brassica oleracea. Phytochemistry 28:1405–1407.

49. Mandel SJ, Shoaf ML, Braco JT, Silver WL, Johnson EC. 2018. Behavioral aversion to AITC requires both painless and dTRPA1 in Drosophila. Front. Neural Circuits 12:45.

50. Mantel N. 1966. Evaluation of survival data and two new rank order statistics arising in its consideration. Cancer Chemother. Rep. 50:163–170.

51. Mariani V, Biasini M, Barbato A, Schwede T. 2013. lDDT: a local superposition-free score for comparing protein structures and models using distance difference tests. Bioinformatics 29:2722–2728.

52. Martin NA. 2004. History of an invader,Scaptomyza flava(fallen, 1823) (Diptera: Drosophilidae). N. Z. J. Zool. 31:27–32.

53. Matsunaga T, Reisenman CE, Goldman-Huertas B, Brand P, Miao K, Suzuki HC, Verster KI, Ramírez SR, Whiteman NK. 2021. Evolution of Olfactory Receptors Tuned to Mustard Oils in Herbivorous Drosophilidae. Mol. Biol. Evol. 39:msab362.

54. Mirdita M, Schütze K, Moriwaki Y, Heo L, Ovchinnikov S, Steinegger M. 2022. ColabFold: making protein folding accessible to all. Nat. Methods 19:679–682.

55. Mithöfer A, Boland W. 2012. Plant defense against herbivores: chemical aspects. Annu. Rev. Plant Biol. 63:431–450.

56. Nishimura Y, Yamada K, Okazaki Y, Ogata H. 2024. DiGAlign: Versatile and interactive visualization of sequence alignment for comparative genomics. Microbes Environ. 39:ME23061.

57. Noge K, Becerra JX. 2015. 4-Oxo-(E)-2-hexenal produced by Heteroptera induces permanent locomotive impairment in crickets that correlates with free thiol depletion. FEBS Open Bio 5:319–324.

58. Oh SM, Jeong K, Seo JT, Moon SJ. 2021. Multisensory interactions regulate feeding behavior in Drosophila. Proc. Natl. Acad. Sci. U. S. A. 118:e2004523118.

59. Orre GUS, Wratten SD, Jonsson M, Hale RJ. 2010. Effects of an herbivore-induced plant volatile on arthropods from three trophic levels in brassicas. Biol. Control 53:62–67.

60. Peláez JN, Gloss AD, Goldman-Huertas B, Kim B, Lapoint RT, Pimentel-Solorio G, Verster KI, Aguilar JM, Nelson Dittrich AC, Singhal M, et al. 2023. Evolution of chemosensory and detoxification gene families across herbivorous Drosophilidae. G3 [Internet] 13. Available from: 10.1093/g3journal/jkad133

61. Peláez JN, Gloss AD, Ray JF, Chaturvedi S, Haji D, Charboneau JLM, Verster KI, Whiteman NK. 2022. Evolution and genomic basis of the plant-penetrating ovipositor: a key morphological trait in herbivorous Drosophilidae. Proc. Biol. Sci. 289:20221938.

62. Rambaut A. 2009. FigTree, version 1.4.3. *Computer program distributed by the author, website:* http://tree.bio.ed.ac.uk/software/figtree/ [accessed January 4, 2011] [Internet]. Available from: https://scholar.google.com/citations?view_op=view_citation&hl=en&citation_for_view=JiYPDfoAAAAJ:W5xh706n7nkC

63. Ray A, van Naters W, van der G, Shiraiwa T, Carlson JR. 2007. Mechanisms of odor receptor gene choice in Drosophila. Neuron 53:353–369.

64. Reichstein T, von Euw J, Parsons JA, Rothschild M. 1968. Heart poisons in the monarch butterfly. Some aposematic butterflies obtain protection from cardenolides present in their food plants. Science 161:861–866.

65. Reisenman CE, Scott K. 2019. Food-derived volatiles enhance consumption in Drosophila melanogaster. J. Exp. Biol. 222:jeb202762.

66. Scott K. 2018. Gustatory processing in Drosophila melanogaster. Annu. Rev. Entomol. 63:15–30.

67. Semmelhack JL, Wang JW. 2009. Select Drosophila glomeruli mediate innate olfactory attraction and aversion. Nature 459:218–223.

68. Shiraiwa T. 2008. Multimodal chemosensory integration through the maxillary palp in Drosophila. PLoS One 3:e2191.

69. Southwood T. 1972. insect/plant relationship--an evolutionary perspective. Available from: https://www.semanticscholar.org/paper/insect%2Fplant-relationship--an-evolutionary-Southwood/635ed400c02b62dca75e90d7ee3e664383d480be

70. Stamatakis A. 2014. RAxML version 8: a tool for phylogenetic analysis and post-analysis of large phylogenies. Bioinformatics 30:1312–1313.

71. Stensmyr MC, Dweck HKM, Farhan A, Ibba I, Strutz A, Mukunda L, Linz J, Grabe V, Steck K, Lavista-Llanos S, et al. 2012. A conserved dedicated olfactory circuit for detecting harmful microbes in Drosophila. Cell 151:1345–1357.

72. Takagi S, Sancer G, Abuin L, Stupski SD, Roman Arguello J, Prieto-Godino LL, Stern DL, Cruchet S, Álvarez-Ocaña R, Wienecke CFR, et al. 2024. Olfactory sensory neuron population expansions influence projection neuron adaptation and enhance odour tracking. Nat. Commun. 15:7041.

73. Tocmo R, Veenstra JP, Huang Y, Johnson JJ. 2021. Covalent modification of proteins by plant-derived natural products: Proteomic approaches and biological impacts. Proteomics 21:e1900386.

74. Wang Y, Qiu L, Wang Bing, Guan Z, Dong Z, Zhang J, Cao S, Yang L, Wang Bo, Gong Z, et al. 2024. Structural basis for odorant recognition of the insect odorant receptor OR-Orco heterocomplex. Science 384:1453–1460.

75. War AR, Paulraj MG, Ahmad T, Buhroo AA, Hussain B, Ignacimuthu S, Sharma HC. 2012. Mechanisms of plant defense against insect herbivores. Plant Signal. Behav. 7:1306–1320.

76. Winde I, Wittstock U. 2011. Insect herbivore counteradaptations to the plant glucosinolate-myrosinase system. Phytochemistry 72:1566–1575.

77. Wu X, Huang H, Childs H, Wu Y, Yu L, Pehrsson PR. 2021. Glucosinolates in Brassica vegetables: Characterization and factors that influence distribution, content, and intake. Annu. Rev. Food Sci. Technol. 12:485–511.

78. Yaffe PB, Power Coombs MR, Doucette CD, Walsh M, Hoskin DW. 2015. Piperine, an alkaloid from black pepper, inhibits growth of human colon cancer cells via G1 arrest and apoptosis triggered by endoplasmic reticulum stress: PIPERINE INHIBITS COLON CANCER CELL GROWTH. Mol. Carcinog. 54:1070–1085.

79. Yao CA, Ignell R, Carlson JR. 2005. Chemosensory coding by neurons in the coeloconic sensilla of the Drosophila antenna. J. Neurosci. 25:8359–8367.

80. Zhao J, Chen AQ, Ryu J, Del Mármol J. 2024. Structural basis of odor sensing by insect heteromeric odorant receptors. Science 384:1460–1467.

